# Directional selection limits ecological diversification and promotes ecological tinkering during the competition for substitutable resources

**DOI:** 10.1101/292821

**Authors:** Benjamin H. Good, Stephen Martis, Oskar Hallatschek

## Abstract

Microbial communities can evade competitive exclusion by diversifying into distinct ecological niches. This spontaneous diversification often occurs amid a backdrop of directional selection on other microbial traits, where competitive exclusion would normally apply. Yet despite their empirical relevance, little is known about how diversification and directional selection combine to determine the ecological and evolutionary dynamics within a community. To address this gap, we introduce a simple, empirically motivated model of eco-evolutionary feedback based on the competition for substitutable resources. Individuals acquire heritable mutations that alter resource uptake rates, either by shifting metabolic effort between resources or by increasing overall fitness. While these constitutively beneficial mutations are trivially favored to invade, we show that the accumulated fitness differences can dramatically influence the ecological structure and evolutionary dynamics that emerge within the community. Competition between ecological diversification and ongoing fitness evolution leads to a state of diversification-selection balance, in which the number of extant ecotypes can be pinned below the maximum capacity of the ecosystem, while the ecotype frequencies and genealogies are constantly in flux. Interestingly, we find that fitness differences generate emergent selection pressures to shift metabolic effort toward resources with lower effective competition, even in saturated ecosystems. We argue that similar dynamical features should emerge in a wide range of models with a mixture of directional and diversifying selection.

---

Ecological diversification and competitive exclusion are opposing evolutionary forces. Conventional wisdom suggests that most new mutations are subject to competitive exclusion, while ecological diversification occurs only under highly specialized conditions (1). Recent empirical evidence from microbial, plant, and animal populations has started to challenge this assumption, suggesting that the breakdown of competitive exclusion is a more common and malleable process than is often assumed (2, 3, 4). Particularly striking examples have been observed in laboratory evolution experiments, in which primitive forms of ecology evolve from a single ancestor over years (5), months (6), and even days (7).

In the simplest cases, the population splits into a pair of lineages, or *ecotypes*, that stably coexist with each other due to frequency-dependent selection, leading to a breakdown of competitive exclusion (5, 6, 8, 9, 10). But evolution does not cease after ecological diversification occurs. Instead, recent sequencing studies have shown that adaptive mutations continue to accumulate within each ecotype, even when population-wide fixations are rare (11, 12, 13). This additional evolution can cause the ecological equilibrium to wander over longer timescales, as observed in the shifting population frequencies of the two ecotypes (5, 13). In certain cases, these evolutionary perturbations can even drive one of the original lineages to extinction, either through the outright elimination of the niche (9), or by the invasion of individuals that mutate from the opposing ecotype (12).

Pairwise coexistence is the simplest form of community structure, but similar dynamics have been observed in more complex communities as well. Some laboratory experiments diversify into three or more ecotypes (7, 14, 15), and it is likely that previously undetected ecotypes may be present in existing experiments (13). Moreover, many natural microbial populations evolve in communities with tens or hundreds of ecotypes engaged in various degrees of competition and coexistence (16, 17, 18). Although the evolutionary dynamics within these communities are less well-characterized, recent work suggests that similar short-term evolutionary processes can occur in these natural populations as well (19, 20, 21).

While the interactions between microbial adaptation and ecology are known to be important empirically, our theoretical understanding of this process remains limited in comparison. Early work in the field of adaptive dynamics (22) showed how ecological diversification emerges under very general models of frequency-dependent trait evolution, which are thought to describe the limiting behavior of a wide class of ecological interactions near the point of diversification. Numerous studies have also investigated the effects of evolution on ecological diversification and stability using computer simulations, in which the parameters of a particular ecological model are allowed to evolve over time (23, 24, 25, 26, 27, 28, 29). However, while both approaches can reproduce some of the qualitative behaviors observed in experiments, it has been difficult to forge a more quantitative connection between these models and the large amount of molecular data that is now available.

One of the main reasons why quantitative comparisons have been difficult is that in many microbial populations of interest, natural selection acts on a range of traits in addition to those directly involved in diversification. The mutations that influence ecological phenotypes must therefore compete with constitutively beneficial mutations at unrelated loci, which can often comprise the bulk of the observed mutation events (12, 13). Although many models exist for describing these constitutively beneficial (or deleterious) mutations in the absence of ecology (30), we have few quantitative predictions for how they behave when they are linked to ecological phenotypes, and vice versa. This makes it difficult to draw any quantitative inferences from the vast molecular data that is now available.

To start to bridge this gap, we introduce a simple, empirically motivated model that describes the interplay between ecological diversification and directional selection at a large number of linked loci. The ecological interactions derive from a well-studied class of consumer resource models (31, 32, 33, 34), in which individuals compete for multiple substitutable resources (e.g. different carbon sources) in a well-mixed environment. We extend this ecological model to allow for heritable mutations in resource uptake rates, which can either divert metabolic effort between resources, or the increase overall fitness. The latter class of mutations provides a natural way to model adaptation at linked genomic loci.

These constitutively beneficial mutations might seem like an ecologically trivial addition to the model, since they are always favored to invade on short timescales. On longer timescales, however, we will show that these accumulated fitness differences can dramatically influence both the ecological structure and the evolutionary dynamics that take place within the community. By focusing on the weak mutation limit, we derive analytical expressions for these dynamics in the two resource case, and we show how our results extend to larger communities as well. These analytical results provide a general framework for integrating ecological and population-genetic processes in evolving microbial communities, and suggest new ways in which these processes might be inferred from time-resolved molecular data.

## EVOLUTIONARY MODEL OF RESOURCE COMPETITION

To investigate the interactions between ecological diversification and directional selection, we focus on a simple ecological model in which individuals compete for an assortment of externally supplied resources in a well-mixed, chemostat-like environment (Fig. 1). This resource-based model aims to capture some of the key ecological features observed in certain microbial evolution experiments (5, 8), as well as more complex ecosystems such as the gut microbiome (18), while remaining as analytically tractable as possible.

**FIG. 1.**
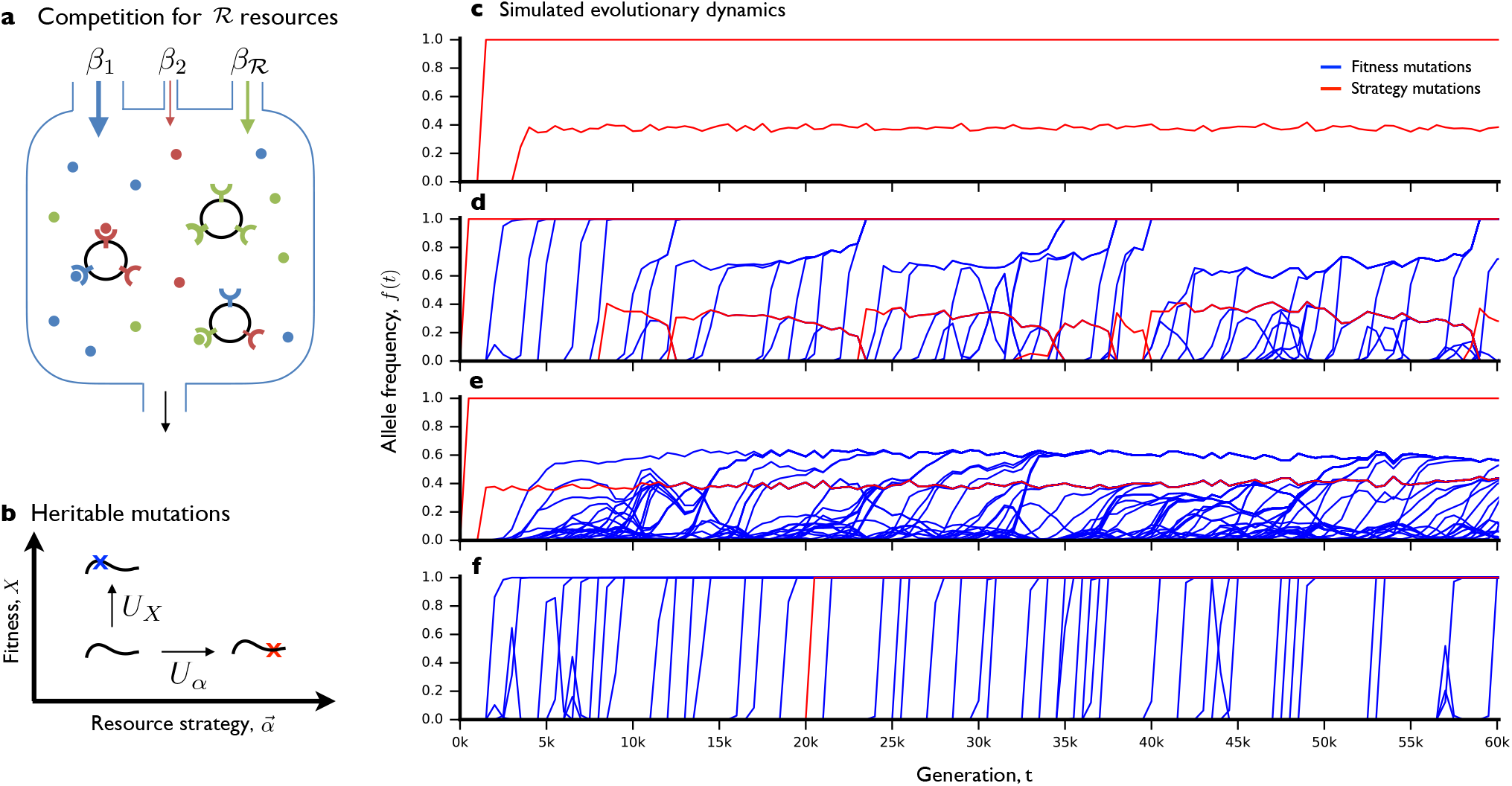
Ecological and evolutionary dynamics in a simplified consumer-resource model. (a) Schematic depiction of ecological dynamics. Substitutable resources are supplied to the chemostat at constant rates 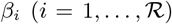, measured in units of biomass 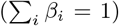. Cells import resources at genetically encoded rates, *r_i_*, which define a normalized *resource strategy* 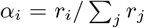 and *general fitness* 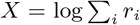. (b) Schematic depiction of evolutionary dynamics. Mutations that alter resource strategies 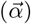 occur at rate *U_α_*, while mutations that alter general fitness (*X*) occur at rate *U_X_*. (c-f) Simulated ecological and evolutionary dynamics, starting from a clonal ancestor, in an environment with 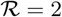 resources. The four panels represent independent populations evolved under different sets of parameters, which differ only in the mutation rates and fitness benefits of pure fitness mutations (SI Appendix E). Lines denote the population frequency trajectories of all mutations that reached fequency ≥ 10% in at least one timepoint. Resource strategy mutations are shown in red, while pure fitness mutations are shown in blue.

In our idealized setting, individuals compete for 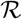 substitutable resources, which are supplied by the environment at a fixed rates (Fig. 1). Individuals are characterized by a resource utilization vector 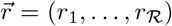, which describes how well they can grow on each of the resources. We assume that the resource utilization phenotypes are constitutively expressed, so that we may neglect complicating factors like regulation. We will find it useful to decompose the phenotype 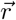 into a normalized portion 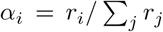, and an overall magnitude 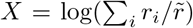, where 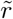 is an arbitrary dimensionful constant. The components of 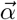 can be interpreted as the fractional effort devoted to growth on resource i, and we will refer to this quantity as the *resource strategy vector*. In contrast, the magnitude *X* resembles an environment-independent measure of *general fitness*, an analogy that we will make more precise below.

We assume that individuals reproduce asexually, so that the state of the ecosystem can be described by the number of individuals *n_μ_* with a given resource strategy vector 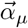 and fitness *X_μ_*. Under suitable assumptions, the ecosystem can be described by the coarse-grained Langevin dynamics,

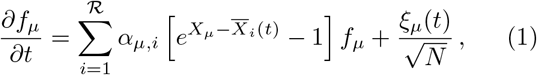

where *N* is a fixed carrying capacity, *f_μ_* = *n_μ_*/*N* is the relative frequency of strain *μ*, and *ξ_μ_*(*t*) is a stochastic noise term (SI Appendix A). The state of the environment is encoded by the set of resource-specific mean fitnesses,

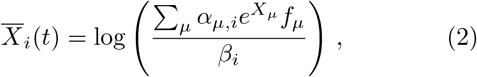

where *β_i_* denotes the fractional flux supplied by resource *i*. Eq. (1) is an example of a more general and well-studied class of consumer resource models introduced by Refs. (31, 35), whose ecological properties have been explored in several recent works (32, 33, 34). The same equation also describes a “bet-hedging” scenario in which the population is periodically subdivided into different environments (SI Appendix A). For a single resource, Eq. (1) reduces to the standard Wright-Fisher model of population genetics (36), with its logistic growth function, *∂_t_* log 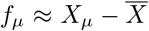. The parameter *X_μ_* can therefore be identified with the (log) fitness in these models. In a multi-resource environment, the deterministic dynamics become more complex, and do not generally admit a closed form solution for *f_μ_*(*t*). However, Eq. (1) still possesses a convex Lyapunov function (SI Appendix A), which implies that *f_μ_*(*t*) must reach a unique and stable equilibrium at long times.

The ecological model in Eq. (1) describes the competition between a fixed set of strains. To incorporate evolution, we also allow for new strains to be created through the process of mutation. We will show that it is useful to distinguish between two broad classes of mutations. The first class, which we will refer to as *strategy mutations*, alter the resource uptake strategy 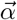, while for simplicity, leave the general fitness *X* unchanged. We assume that these mutations occur at a per genome rate *U_α_* and result in a new resource strategy 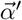 drawn from some distribution 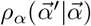. In addition to these strategy mutations, we consider a second class of *pure fitness mutations*, which alter the general fitness *X* but leave the resource strategy 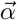 unchanged. These mutations capture the effects of directional selection at a large number of other loci throughout the genome, which may only be tangentially related to the resource utilization strategy. We assume that these fitness mutations arise at a per-genome rate *U_X_*, and that they increment *X* by an amount *s* drawn from the distribution of fitness effects, *ρX*(*s*). For simplicity, we assume that there is no macrosopic epistasis for fitness (37), so that *ρX*(*s*) remains the same for all genetic backgrounds.

This division into fitness and strategy mutations is neither exhaustive nor unambiguous. Some changes in resource strategy may also incur a fitness cost, and one can simulate a pure fitness mutation by shifting metabolic effort away from resources that are not present in the current environment (i.e., those with *β_i_* =0). Nevertheless, by considering these as independent axes, we will show that we can gain additional insight into the behavior of our model.

Pure fitness mutations might seem like an ecologically trivial addition to the model, because they are always favored to invade. However, computer simulations show that these accumulated fitness differences can still have a dramatic influence on both the ecological structure and the evolutionary dynamics that arise in a given population. Figures 1C-F depict individual-based simulations of four populations, which are subject to the same environmental conditions and the same supply of strategy mutations, but have different values of *U_X_* and *ρX*(*s*) (SI Appendix E). Depending on the supply of fitness mutations, the behaviors range from rapid diversification and stasis (Fig. 1C) to unstable but continually renewed coexistence (Fig. 1D), stable coexistence and rapid within-clade evolution (Fig. 1E), and the permanent disruption of coexistence (Fig. 1F).

To understand these different behaviors and their dependence on the underlying parameters, we will start by analyzing the simplest non-trivial scenario, in which the strains evolve in an environment with just two resources. In this case, the environmental supply rates and resource uptake strategies can be described by scalar parameters 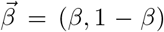 and 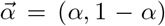, respectively. This case will already be sufficient to elucidate many of the key qualitative behaviors and fundamental timescales involved, while maximizing analytical tractability. In the second section, we will extend this analysis to larger numbers of resources, and comment on the additional features that arise only in this more complicated scenario.

## ANALYSIS

### Selection for ecosystem to match environment, stable coexistence

We will begin by considering the dynamics in the absence of fitness differences (*U_X_* = 0, *X_μ_* = 0). The ecological dynamics in this “neutral” scenario have recently been described by Ref. (32), and it will be useful to build on these results in the sections that follow.

We begin by considering a single strategy mutation that occurs in a clonal population of type *α*_1_, creating a new strain of type *α*_2_. The initial dynamics of this mutation can be described by a branching process with growth rate *S*_inv_ = 〈*∂_t_f*〉/*f* (SI Appendix B), also known as the *invasion fitness*. In this case, the invasion fitness is given by

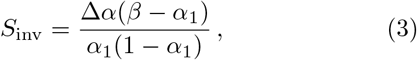

where Δ*α* = *α*_2_ – *α*_1_ is the difference between the mutant and wildtype uptake rates. The invasion fitness is positive whenever Δ*α* and *β* – *α*_1_ have the same sign: if *α*_1_ < *β*, then selection will favor mutations that increase *α*, while if *α*_1_ > *β*, selection will favor mutations that decrease *α*. In this way, selection tries to tune the population uptake rate to match the environmental supply rate. If *α*_1_ = *β*, then the invasion fitness vanishes for all further strategy mutants. This constitutes a marginal evolutionarily stable state (ESS). It will turn out that many expressions simplify in a *near-ESS limit*, where the uptake rates *αi* remain close to *β*. We will focus on this regime for the expressions quoted in the remainder of the main text, while the full expressions are derived in SI Appendix.

Since all mutations start as a single copy in the population, many will be lost due to genetic drift, even when they are favored by selection. With probability ~*S*_inv_, the mutant lineage will survive to reach frequency *f* ~ 1/*NS*_inv_, and will then start to increase deterministically; in large populations, this will typically occur long before the mutant starts to influence its own growth rate, so that the constant invasion fitness assumption is justified (SI Appendix B).

At long times, the ecological dynamics will lead to one of two final states: the mutant will either replace the wildtype (competitive exclusion) or the two will coexist at some intermediate frequency (Fig. 2A). The latter scenario will occur if and only if the wildtype can re-invade a population of mutants, which requires that the reciprocal invasion fitness, 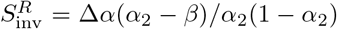, is also positive. By examining this expression, we see that the mutant will outcompete the wildtype if its strategy lies between *β* and *α*_1_, while stable coexistence occurs when *α*_1_ and *α*_2_ span *β* (i.e., *α*_1_ < *β* < *α*_2_ or vice versa). When this condition for coexistence is met, Ref. (32) has shown that the steady-state frequencies are determined by the linear equation,

**FIG. 2.**
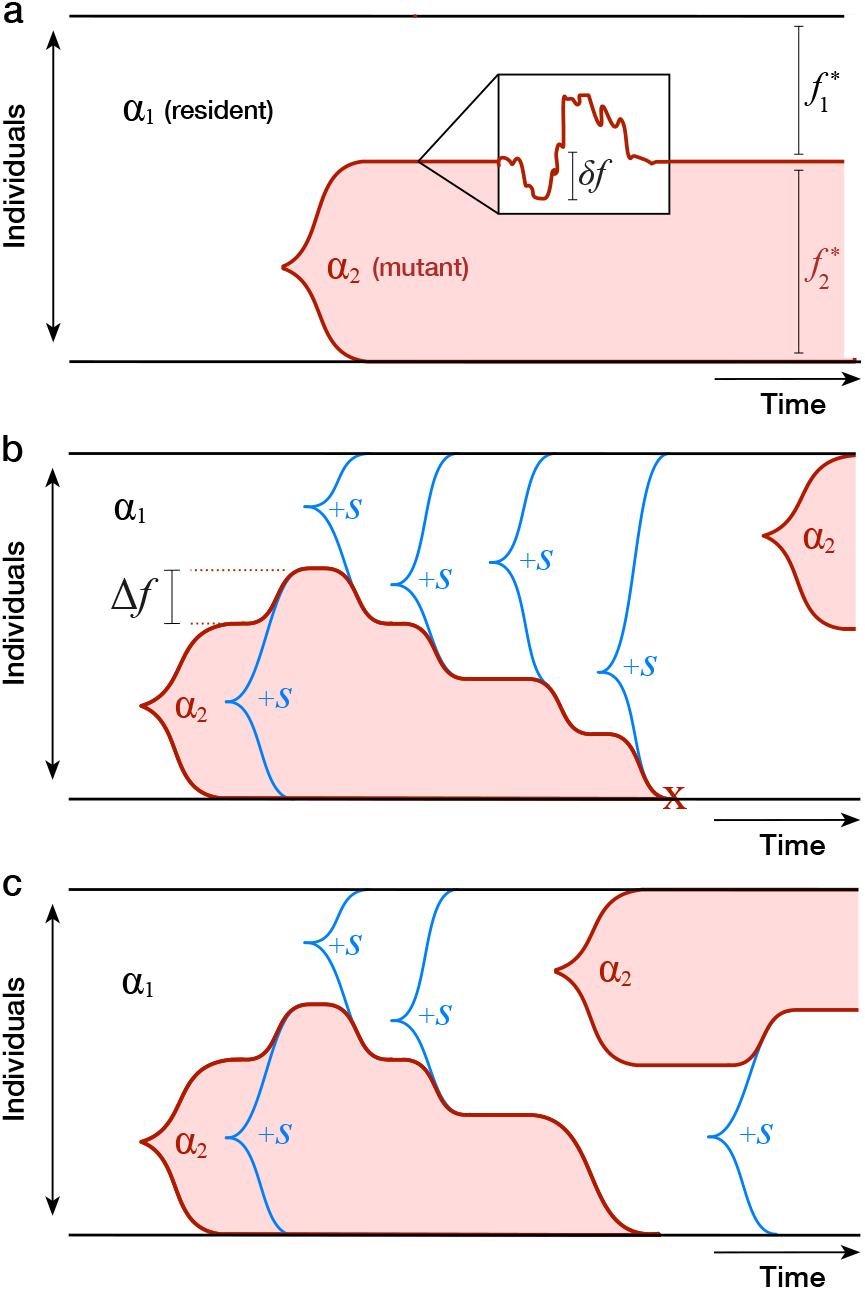
Schematic illustration of key eco-evolutionary processes in a two-resource ecosystem. (a) Ecological diversification from a clonal ancestor. In the absence of fitness mutations, strains coexist at a stable equilibrium (*f*\*) with fluctuations (*δf*) controlled by genetic drift. Further strategy mutations are not favored to invade. (b) Pure fitness mutations that sweep within an ecotype shift the stable equilibrium by Δ*f*; accumulated fitness differences can ultimately drive ecotypes to extinction. Further strategy mutations allow the winning clade to re-diversify at a later time. (c) Occupied niches can also be invaded by strategy mutations that arise in fitter genetic backgrounds. In this case, the original ecotype lineage is driven to extinction while the ecological structure of the community is preserved.

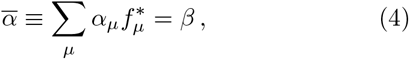

whose solution is given by *f*\*/(1–*f*\*) = (*β*–*α*_2_)/(*α*_1_ – *β*). In other words, the relative frequencies of the strains are inversely proportional to their distance from the environmental supply rate. According to Eq. (4), these frequencies are chosen such that the population-averaged uptake rate 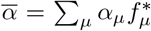 exactly balances the resource supply rate *β*. This provides an intuitive explanation for the cause of coexistence: by maintaining the strains at intermediate frequencies, the population is able to match the environmental supply rate more closely than it could with either strain on its own.

Once this ecological equilibrium is attained, number fluctuations will continuously perturb the true frequency away from *f*\* (Fig. 2A), subject to a linearized restoring fitness ~Δ*α*^2^/*β*(1 – *β*) (SI Appendix B). The restoring force is strong compared to genetic drift when *N*Δ*α*^2^/*β*(1 – *β*) ≫ 1, which leads to linearized fluctuations of order 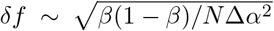, and a lifetime for the stable state that is exponentially long in 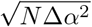. At this point, additional strategy mutants are subject to very weak selection pressures: fluctuations will induce momentary invasion fitnesses of order 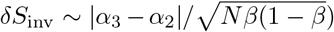 (which can be large compared to 1/*N*), but these fitness effects are quickly averaged to zero during the ~1/*δS*_inv_ generations required for such a mutation to establish (SI Appendix B). Thus, once the population diversifies to fill the two niches, the rate of evolution dramatically slows down (as, e.g. in Fig. 1A), since the relevant timescales are controlled by genetic drift. In this way, a large effect mutation can allow the ecosystem as a whole to reach an effective ESS, long before any of the constituent strains reach the ESS on their own.

### Diversification load

We are now in a position to analyze how fitness alters the basic picture above. We begin by revisiting the invasion of a mutant strain in an initially clonal population, this time allowing for a fitness difference Δ*X* between the mutant and wildtype. In this case, the new invasion fitness is given by a simple linear combination,

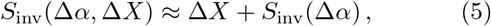

where *S*_inv_(Δ*α*) is the invasion fitness for a pure strategy mutation from Eq. (3). This result describes, in quantitative terms, how selection balances its ecological preferences 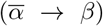 with its desire to maximize fitness 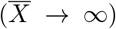. When the uptake rate of the resident population is far from the environment supply rate 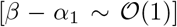, the ecological selection pressures can be quite strong, with invasion fitnesses as high as 10% – 100%. This implies that strongly deleterious mutations of order

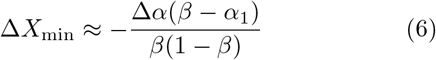

can hitchhike to fixation when the population colonizes a new ecological niche (a form of *diversification load*).

### Fitness differences perturb ecological equilibria

In addition to shifting the invasion fitness of a new mutation, fitness differences can also alter the long-term ecological equilibrium between mutant and wildtype in Eq. (4). In the extreme limit, this can disrupt the stable coexistence altogether. If the mutant is less fit than the wildtype (Δ*X* < 0), this will occur whenever Δ*X* is less than the maximum diversification load Δ*X*_min_ in Eq. (6). On the other hand, if Δ*X* > 0, extinction will occur when the wildtype no longer has positive invasion fitness, or when Δ*X* exceeds a threshold

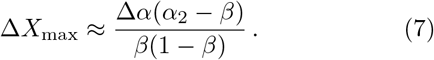

We note that the fitness differences in Eqs. (6) and (7) are lower than the values required for the mutant or wild-type to dominate in *all* environmental conditions (SI Appendix B). Instead, the fitness thresholds strongly depend on how the resource strategies differ from each other, and from the environmental supply rate. When Δ*α* ~ *ϵ*, even a small fitness difference (Δ*X* ~ *ϵ*^2^) can disrupt the stable ecology, while for 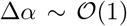, much larger fitness differences (Δ*X* ≳ 100%) can be tolerated.

When Δ*X*_min_ > Δ*X* < Δ*X*_max_, the two strains continue to coexist, but their equilibrium frequency is no longer given by Eq. (4). In this case, the competing drive to maximize fitness means that selection will no longer favor an ecology that matches the environmental resource distribution, at least not perfectly. In SI Appendix B, we show that the new equilibrium frequency is given by

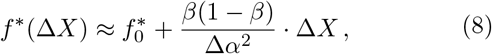

where 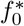 is the neutral ecological equilibrium from Eq. (4). From this expression, we can read off the typical fitness differences required to perturb *f*\* from its present value. This fitness sensitivity is again determined by the distance between the two resource strategies. If 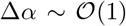, large fitness differences (Δ*X* ≳ 100%) are required to change the equilibrium frequency, while for Δ*α* ~ *ϵ*, even very small fitness differences (Δ*X* ~ *ϵ*^2^) can generate large changes in the equilibrium frequency.

### Further fitness evolution and diversification-selection balance

Once the population achieves the stable ecology in Eq. (8), additional fitness mutations will occur in each strain with probability proportional to the equilibrium frequency *f*\*. In our model, the invasion fitness of such a mutation is simply its fitness effect *s*, independent of the ecological state of the population. With probability ~*s*, this mutation will sweep through its parent clade, changing the fitness difference between the clades by ±*s* and the equilibrium frequency by Δ*f* = *f*\*(Δ*X*±*s*) –*f*\*(Δ*X*) (Fig. 2B). In the linear regime of Eq. (8), the frequency and fitness changes are directly related,

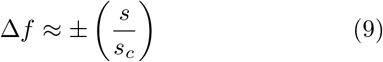

where *s_c_* = Δ*α*^2^/*β*(1 – *β*) is the fitness scale that determines changes in equilibrium frequency. If *s* ≫ *s_c_f*\*(1 – *f*\*), then the stable coexistence will be di srupted, and the mutant clade will take over the population. We will refer to such a scenario as *ecosystem collapse*, since one of the niches is no longer occupied.

Similar behavior can occur when *s* ≪ *s_c_* as well, except that now the ecosystem collapse occurs due to cumulative effect of many general fitness mutations. When the fitness mutations accumulate independently, this process can be described by an effective diffusion model,

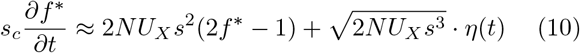

with a bias that reflects the higher probability of producing a mutation in a larger clade (SI Appendix B). Eq. (10) superficially resembles the drift-induced perturbations at ecological equilibrium, except that the bias is now unstable rather than restoring. When 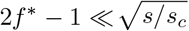, the mutation bias is weak, and the clade frequencies undergo a random walk 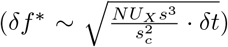. But after a time of order 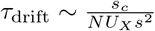, the frequency differential grows large enough that the more prevalent clade will deterministically produce more beneficial mutations, so that it is destined for fixation. After a time of order 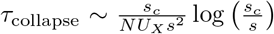, the fitness difference between the clades grows so large that the ecosystem finally collapses (Fig. 2B). This timescale sets an upper limit on the lifetime of the stable state when many fitness mutations are available.

Once the ecosystem collapses, there will be a strong selection pressure for the winning clade to re-diversify through additional strategy mutations, and restart this process from the beginning (Fig. 2B). To gain insight into these dynamics, we first consider the case where the resource strategies are controlled by a single genetic locus, with fixed phenotypes *α*_1_ and *α*_2_, and mutations that alternate between the two states at rate *U_α_*. After an ecosystem collapse, Eq. (3) shows that the invasion fitness for the opposite strategy is given by *S*_inv_ ~ *s_c_*, so the collapsed state will persist for a time of order *τ*_diversify_ ~ 1/*NU_α_s_c_*, until the stable ecology is reestablished. If the two strategies are symmetric about *β*, so that *f*\*(0) = 1/2, the new stable state will persist for ~*τ*_collapse_ generations in the absence of additional strategy mutations, and the process will then repeat itself. The relative probability of observing the population in the collapsed 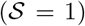 or saturated 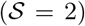 states is therefore given by

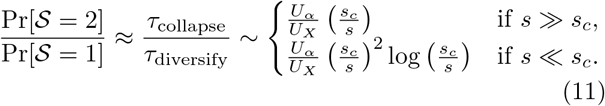

This expression shows the minimum amount of strategy mutations, or the maximum amount of general fitness mutations, that allow the population to maintain a saturated ecosystem. We will refer to this dynamic steady state as *diversification-selection balance*, in analogy to mutation-selection balance in population genetics (38). Note that this balance crucially depends on the state of the ecosystem through *s_c_* ~ Δ*α*^2^/*β*(1 – *β*). All else being equal, ecosystems with more similar resource uptake strategies will be disrupted more easily than those with a higher degree of specialization.

### Invading ecotypes can delay ecosystem collapse

Strictly speaking, our derivation of Eq. (11) is only valid in the limit that *τ*_collapse_ ≪ *τ*_diversify_, since we neglected mutations between *α*_1_ and *α*_2_ when both niches were filled. When *τ*_collapse_ ≳ *τ*_diversify_ (i.e., when the ecosystem spends an appreciable amount of time in the saturated state), we must also account for mutations between the two strategies that arise before the ecosystem collapses. Those mutations that arise in the less-fit clade will have little chance of invading. However, a mutation from the more-fit to the less-fit strategy will establish with probability ~|Δ*X*|, where Δ*X* is the current fitness difference between the two clades. If this mutation is successful, it will outcompete the resident lineage with the corresponding value of a, and reset the fitness difference to Δ*X* = 0 (Fig. 2C). In this way, invasion from one ecotype to another can significantly delay the process of ecosystem collapse, since it relieves the tension between fitness maximization and 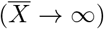 and selection to match the environment 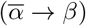.

To analyze this process, we note that successful invasion events will occur as an inhomogeneous poisson process with rate 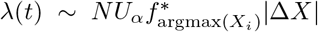, where *f*\*(*t*) and Δ*X*(*t*) are again determined by the diffusion model in Eq. (10). This leads to a characteristic invasion timescale

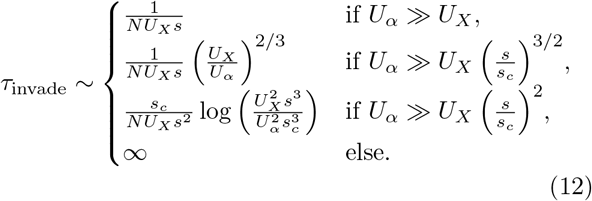

which is derived in SI Appendix B. Each of these regimes corresponds to a different intuitive picture of the dynamics. In the first case, strategy mutations are frequent compared to general fitness mutations, and invasion occurs almost immediately after the first fitness mutation arises. In the second case, invasion occurs after multiple fitness mutations have accumulated, but when the frequencies of the clades still wander diffusively relative to each other 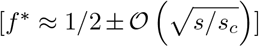. In the third regime, invasion occurs after one of the clades has grown to a sufficiently large frequency that it would have deterministically led to ecosystem collapse. When the invading mutant establishes, it will therefore cause a rapid shift in the frequencies of the ecotypes as *f*\*(Δ*X*) returns to *f*\*(0). _2_

Finally, when 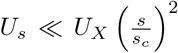, strategy mutants are sufficiently rare that the ecosystem will typically collapse and re-diversify before invasion can occur. This sets the region of validity of the diversification-selection balance in Eq. (11). Interestingly, Eq. (11) shows that collapse and re-diversification can still dominate over invasion even when both niches are typically filled 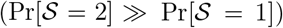. In this case, both the genealogical structure and the typical state of the ecosystem will resemble the invasion regime, but the historical record would contain a series of puncutated extinction and diversification events, interspersed with long periods of gradual fitness evolution.

### Fitness differences create opportunities for ecological tinkering

Our derivation of Eqs. (10) and (12) assumed that the two ecotypes were fixed by the genetic architecture of the organism. Individuals could mutate between *α*_1_ and *α*_2_, but mutations to other points in strategy space were forbidden. In the absence of fitness differences, we saw that selection for these additional strategy mutants is weak once both niches have been filled (*S*_inv_ ≲ 1/*N*), potentially justifying the single-locus assumption in terms of a priority effect. However, the previous analysis shows that there can be strong selection to switch strategies once fitness mutations accumulate, so it is also plausible that fitness differences could lead to selection for strategy mutations more generally.

To investigate these selection pressures, we consider a population that is currently described by the steady state in Eq. (8). We then consider strategy mutations that occur on the background of *α*_2_, altering its strategy to *α*_3_ while leaving its fitness intact. The invasion fitness for such a mutation is given by

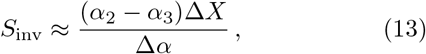

As anticipated, fitness differences create additional selection pressure for strategy mutations beyond the simple switching behavior considered above.

The direction of selection is determined by the sign of Δ*X*. In the background of the fitter clade (Δ*X* > 0), selection favors mutations that increase the strategy in the direction of *β* (a form of generalism), while simultaneously disfavoring mutations that lead to increased specialization (Fig. 3). The opposite behavior occurs in the less-fit background, with selection favoring mutations that increase the distance from *β*, leading to increased specialization. Both behaviors have an intuitive explanation in terms of individuals preferring to allocate their metabolic energy toward the resource with the least-fit consumers, thereby minimizing the effective competition that they experience.

**FIG. 3.**
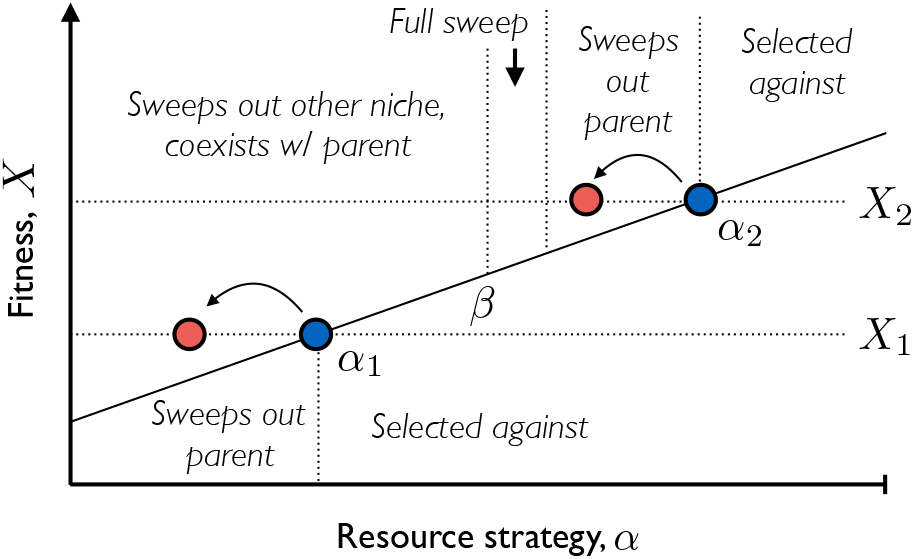
Invasion fitness landscape for additional strategy mutations in a two-resource ecosystem. The two resident ecotypes are illustrated by blue circles, while red circles denote mutant strains created by strategy mutations on one of the ecotype backgrounds. The solid black line denotes the effective mean fitness, 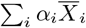, experienced by a given resource strategy. Strains with general fitness (*X*) above this line are favored to invade, while others are selected against. If a mutant successfully invades, its effect on the ecosystem is indicated by the text.

Once a successful strategy mutation arises, it will sweep through the population and alter the ecological equilibrium (Fig. 3). Mutations in the less-fit clade are straightforward to analyze. Since these are always directed away from both *β* and *α*_1_, these mutants will sweep through their parent clade and increase the equilibrium frequency according to Eq. (8). Successful mutations in the fitter clade have a wider range of outcomes, since these are always directed toward *β* and *α*_1_. If *α*_3_ < *β*, the mutant lineage will outcompete the less-fit strain *α*_1_, and will stably coexist with its parent clade *α*_2_ at an equilibrium frequency *f*\* = (*β* – *α*_2_)/(*α*_3_ – *α*_2_). On the other hand, if *β* < *α*_3_ < *α*_2_, the mutant lineage will always sweep through its parent clade *α*_2_. If *α*_3_ is sufficiently close to *α*_2_, this will simply lead to an increase in frequency according to Eq. (8). However, if *α*_3_ is close enough to *β* that Δ*X*_max_(*α*_3_) becomes less than the actual fitness difference, Δ*X*, then the mutant will sweep out both clades and lead to an ecosystem collapse and subsequent re-diversification. Thus, in addition to creating a larger target for invasion events, these additional strategy mutations can also enhance the probability of ecosystem collapse. The balance between these competing tendencies will depend on the genetic architecture of the resource strategies, *ρ_α_*(*α*’|*α*), which is poorly parameterized by existing data. A detailed analysis of the potential regimes will be left for future work.

### Beyond pairwise coexistence

Our previous analysis focused on environments with only two substitutable resources, where at most two strains can coexist at equilibrium. In this case, the structure of the stable ecosystem was simple enough to admit a full analytical solution, which we could use to derive explicit predictions for many evolutionary quantities of interest. However, many microbial communities are found in environments with large numbers of potential resources, and flexible gene pools that allow them to alter their resource uptake rates through horizontal gene transfer (39). It is therefore natural to ask how our results generalize to these more complicated environments as well. A full analysis of this case is beyond the scope of the present work, as there are even fewer constraints on the space of ecological and evolutionary parameters compared to the two resource case. Nevertheless, it is still useful to know whether our qualitative results extend beyond 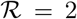, and whether there are fundamentally new behaviors that only arise in higher dimensions.

For a general ecological equilibrium, a mutation that alters the phenotype of a resident strain from 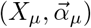 to 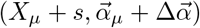 will have an invasion fitness

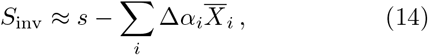

where the resource-specific mean fitnesses 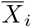 are given by Eq. (2) evaluated at the equilibrium frequencies 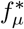 (SI Appendix C.1). Increases in *α_i_* are favored when 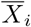 is lower than the “effort-averaged” 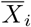 for the other resources, and vice-versa. Thus, similar to the two-resource case, there is still a sense in which selection favors mutations that flow from high values of 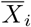 to lower values of 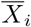, though there are now 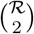 beneficial directions, 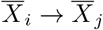, rather than just one.

The invasion fitness in Eq. (14) depends on the current community composition only through the intensive variables 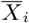. In a *saturated* ecosystem, where the number of coexisting strains is equal to the number of resources, these can be directly obtained by a matrix inversion of Eq. (1),

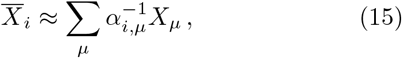

where 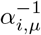 is the left inverse of *α_μ,i_*. Thus, we see that in a saturated ecosystem, the 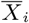 are given by linear combinations of the strain fitnesses *X_μ_*, justifying their interpretation as resource-specific mean fitnesses. Moreover, perturbation expansions of 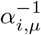 suggest that the prefactor is still inversely proportional to an effective distance between the strategies (SI Appendix C.2), similar to the two-resource case in Eq. (13). We note that the equilibrium values of 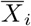 are conditionally independent of both the resource supply vector *β_i_* and the strain frequencies 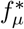; these quantities influence 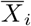 only through shaping the set of resource strategies that coexist at equilibrium. Thus, these saturated ecosystems dynamically adjust their composition to screen the internal selection pressures 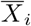 from the external environmental conditions. Similar findings were recently obtained for the neutral case [where 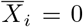 (32)], as well as in certain community assembly processes in the 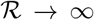 limit (33, 34). Eq. (15) shows that this is a generic property that occurs whenever the number of surviving species is equal to the number of resources.

In this limit, the steady-state frequencies 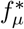 can be obtained from a similar matrix inversion,

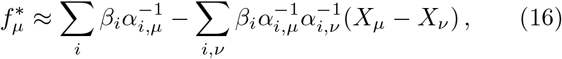

which serves as the generalization of Eq. (8) for 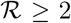 (SI Appendix C.2). As in the two-resource case, small fitness differences perturb the neutral ecological equilibrium via linear combinations of the strain fitnesses, with a prefactor that is inversely proportional to the square of the effective distance between the resource strategies.

While the saturated case is particularly simple, we saw above that fitness mutations can drive the number of surviving species below this saturated value. In contrast to the two-resource case, these *unsaturated* ecosystems can now harbor multiple coexisting strains when 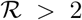, leading to a continuous generalization of the diversification-selection balance in Eq. (11). To investigate this effect, we performed computer simulations of a binary strategy space model, in which individuals can either import or not import a given resource, with mutations that toggle individual uptake rates on and off (SI Appendix C.3). The results recapitulate the qualitative behavior observed for 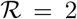 resources, in that a sufficiently high rate of general fitness mutations can constrain the number of distinct strategies that are able to coexist (Fig. 4B). To compensate for the strong ecological selection pressures that can arise when 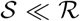, the populations are forced to evolve consortia of “generalist” strains such that 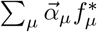 is still close to 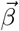, at least at lowest order (Fig. 4A).

**FIG. 4.**
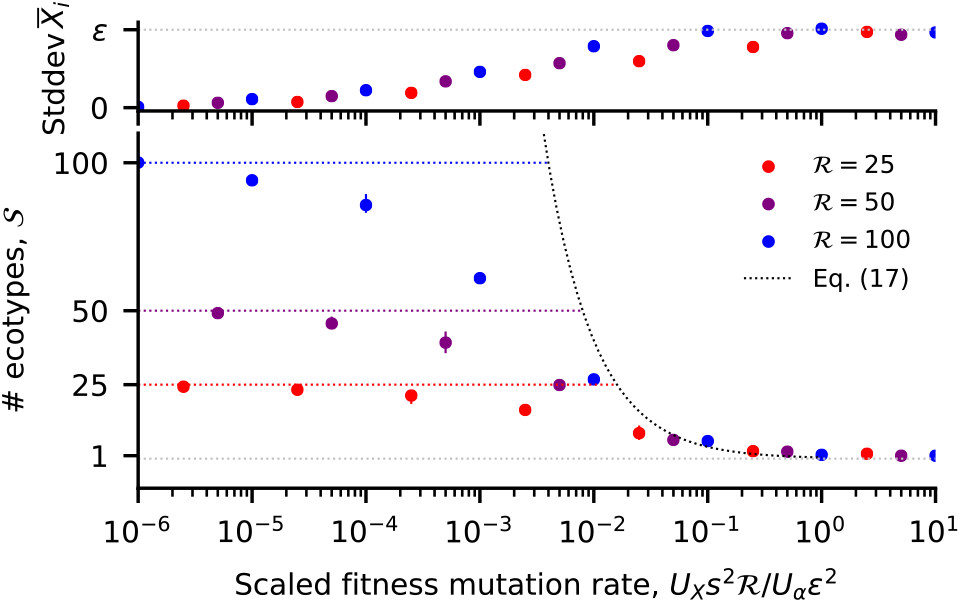
Diversification-selection balance when 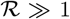. Circles depict the long-term steady state from SSWM simulations of a binary resource usage model in a nearly uniform environment (SI Appendix E.2). Each point denotes an average over multiple timepoints from 3 independent replicates; solid lines indicate the minimum and maximum replicate. (a) The standard deviation in 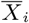 across the 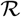 resources. (b) The number of coexisting ecotypes. The colored dashed lines denote the maximum ecosystem capacity 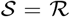. The black dashed line depicts the scaling relation in Eq. (17) which applies for 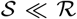, with an 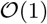 prefactor of 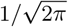 included for visualization.

In a nearly uniform environment 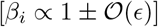, simulations show that the steady-state ecosystem tends to be dominated by a single “generalist” strain (*α_μ,i_* ∝ 1), and a collection of 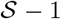 single loss-of-function variants (*α_μ,i_* ∝ 1 – *δ_μ,i_*) that recently descended from mutations in the generalist background (Figs. 5 and S2-S4). These simple community structures can be characterized analytically, and in the weak-mutation limit, yield a simple heuristic expression for the diversification-selection balance,

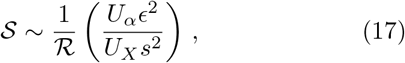

which is valid for 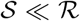 (SI Appendix C.3). The transition to the fully saturated state 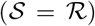 requires an even more stringent condition, which implies that unsaturated ecosystems are obtained for a very broad parameter regime (Fig. 4B). In both cases, a larger number of substitutable resources will lead to a less diverse ecosystem at diversification-selection balance. This is ultimately due to the fact that the difference between generalists and single loss-of-function variants becomes increasingly small as 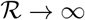.

This suggests that the relative frailty of the diversification-selection balance in Eq. (17) may be a pathological feature of the simple genetic architecture that we have assumed, in which fit generalist phenotypes are easily accessible. If we instead impose an upper limit 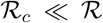 on the number of resources that a strain can metabolize, heuristic calculations suggest that diversification-selection balance will be achieved for substantially higher values of 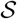, even for large 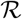 (SI Appendix C.4). In this case, the ecological and genealogical structures that are attained at this evolutionary steady state will be considerably more complex than the shallow star-shaped genealogies in Fig. 5. A more detailed analysis of this steady state will be considered in future work.

**FIG. 5.**
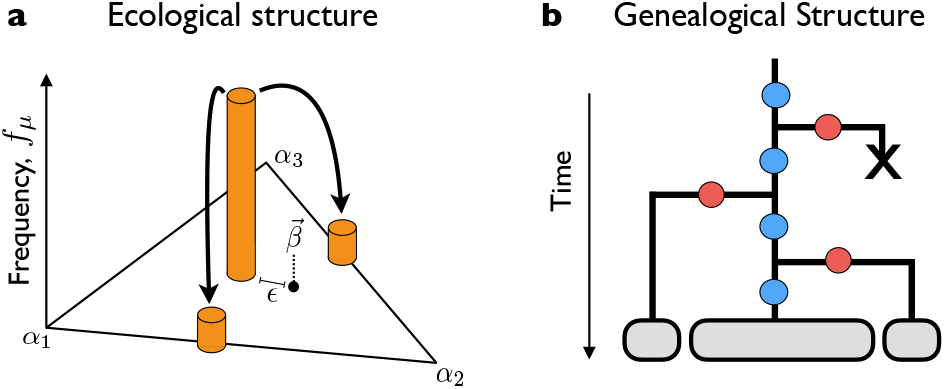
Schematic of (a) ecological and (b) genealogical structure at the evolutionary steady-state described in Eq. (17). In (b), blue dots represent general fitness mutations and red dots represent loss-of-function strategy mutations.

## DISCUSSION

In microbial populations, primitive ecological interactions can evolve spontaneously over years (5), months (6), and even days (7). Yet this process rarely takes place in isolation. In rapidly evolving populations, diversifying selection must compete with directional selection acting on other loci throughout the genome. Here, we have introduced a simple mathematical framework to model the interactions between these two processes in asexually reproducing organisms.

The ecological interactions in our model emerge from the competition for substitutable resources (e.g. different carbon sources), according to a well-studied class of models from theoretical ecology (31, 32, 33, 34). To incorporate evolution into this model, we assumed that individuals can acquire mutations that alter their resource uptake rates. We showed that it is useful to distinguish between two characteristic types of mutations: (i) strategy mutations, which divert metabolic effort from one resource to another and (ii) fitness mutations, which increase the overall growth rate but leave the relative uptake rates unchanged. In this classification scheme, strategy mutations enable ecological diversification, while fitness mutations capture the effects of directional selection at other genomic loci.

The creation of new strains via mutation bears a superficial resemblance to immigration from a fixed species pool, which is the traditional scenario considered in theoretical ecology. However, this analogy is exact only in the absence of inheritance, when the phenotypes of nearby genotypes are uncorrelated from each other. In contrast, when the effects of mutations are heritable, we have seen that directional selection can produce dramatic departures from traditional ecological predictions.

Similar to immigration (34, 40), strategy mutations allow an initially clonal population to diversify into stably coexisting ecotypes, whose upper bound is set by the number of resources. Yet because fitness mutations are heritable, further evolution will lead to fitness differences between the clades, which can dynamically shift the ecological equilibrium over time, and eventually drive less fit clades to extinction. The mere observation that selection can disrupt coexistence is not surprising, since drug resistance or other harsh selection regimes provide striking examples of this effect. However, our quantitative analysis shows that this collapse can happen long before any clade is universally inferior to another, and that it can result from the compound effect of many small effect mutations that would not lead to extinction on their own. These results suggest that ongoing directional selection may have a larger impact on the structure of microbial communities than is often assumed. In particular, while previous ecological analyses suggest that the number of ecotypes should meet (33, 34) or even exceed (32) the number of resources, our results raise the possibility that they could also reside at a *diversification-selection balance* below the maximum capacity of the ecosystem.

In addition to their influence on coexistence, we also found that fitness differences accrued via directional selection will generate emergent selection pressures for continual evolution of the ecological phenotypes, even in a saturated ecosystem. While these internal selection pressures are reminiscent of the Red Queen effect (41), our quantitative analysis shows that they select for different phenotypes than in the standard predator-prey setting. In particular, less fit clades do not experience increased selection pressure to narrow their fitness deficit by accumulating fitness mutations. Instead, selection favors mutations that divert metabolic effort toward resources with lower effective competition, even at the cost of widening the fitness deficit. Moreover, the direction of selection toward any given resource can shift dynamically as the fitness differences and resource uptake strategies evolve over time.

Most of our analysis focused on the strong-selection weak-mutation regime, in which the current ecological equilibrium is attained before the next mutation occurs. In this limit, when the resource uptake strategies are sufficiently close to the supply rates, our model takes on a universal form that closely resembles traditional models of adaptive dynamics (22, 42). The key difference is that directional selection behaves as an additional trait dimension, which is effectively constrained to remain far from its optimum at all times (Fig. S1, SI Appendix D). Our results show that this simple broken symmetry can lead to dramatic deviations from the standard adaptive dynamics picture.

In contrast to adaptive dynamics, we also allow for mutations that have non-infinitesimal effects on resource uptake rates, which turn out to play a key role in controlling the dynamical behaviors that we observe. In practice, the genetic architectures of most ecological interactions remain poorly characterized empirically. In a few well-studied cases, ecological diversification can be traced to a single large-effect mutation (9, 43), while in others, a series of smaller mutations have been implicated (44). Our present analysis suggests new ways in which we might constrain this key parameter experimentally, either by analyzing fluctuations in ecotype frequencies on long timescales (13), or by measuring the joint distribution invasion fitness (*S*_inv_) and ecological perturbation (Δ*f*) across a panel of engineered mutations (44).

Of course, the present work has focused on a highly simplified model, which omits many of the complicating factors expected in either natural or laboratory settings. Future work will be required to fully explore the effects of clonal interference (SI Appendix B.3), time-varying environments, crossfeeding, recombination, and other additions to our basic model. We believe that our results provide a promising analytical framework in which to investigate these effects, which have mostly been confined to simulations so far.

It is also interesting to ask whether our results can mapped onto more diverse modes of ecological interaction, or whether there are other universality classes yet to be discovered. Since our model can be viewed as the simplest generalization of population genetics with multiple fitness axes, we hypothesize that it may capture the limiting behavior of a broad class of ecological interactions that are mediated by a small number of intensive variables. If so, its analytical tractability may offer a promising avenue for investigating the interactions between ecology and evolution more generally.

## ACKNOWLEDGMENTS

BHG thanks Evgeni Frenkel and Ned Wingreen for discussions that inspired the development of the model. BHG acknowledges support from the Miller Institute for Basic Research in Science at the University of California Berkeley. OH acknowledges support from a National Science Foundation Career Award (No. 1555330) and a Simons Investigator award from the Simons Foundation (No. 327934). This research used resources of the National Energy Research Scientific Computing Center, a DOE Office of Science User Facility supported by the Office of Science of the U.S. Department of Energy under Contract No. DE-AC02-05CH11231.

## Appendix A: Derivation of Eqs. (1) and (2)

In this section, we show how the coarse-grained Langevin dynamics in Eqs. (1) and (2) emerge from two different microscopic models. The first is a simplified class of consumer-resource models described in the main text. To illustrate the generality of these dynamics, we also describe a second implementation that is a more direct extension of the traditional Wright-Fisher model, in which subsets of a population are randomly assigned to different environmental conditions.

### 1. Consumer-resource model

Our consumer-resource derivation closely follows the one described by Ref. (32). We assume that all strains *μ* and resources *i* are present in a well-mixed volume *V*, which is diluted at rate *δ*. In the consumer-resource framework, the per capita growth rate of each strain is mediated by the resource concentrations,

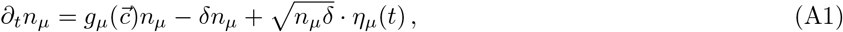

where *n_μ_* is the absolute number of individuals of strain *μ*, 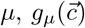 is a strain-specific growth function, and *η_μ_*(*t*) is an uncorrelated Brownian noise term (45). The resource concentrations (in units of *V*^−1^) obey a second set of equations,

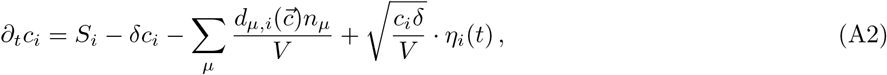

where *S_i_* is the input flux of resource *i* and 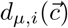 is the per capita depletion rate of resource *i* by strain *μ*. This general class of models has been studied previously by Refs. (31, 35), and others. Following Ref. (32), we consider a restricted subset of models where the growth and depletion functions take on a particularly simple form:

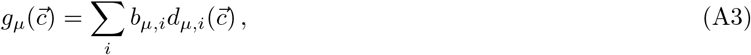

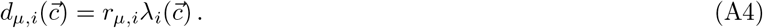

The first assumption states that the resources are *effectively substitutable*, i.e. biomass can be produced equally well from suitably normalized versions of any imported resource. The constant normalization factor 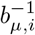 can be interpreted as the amount of imported resource *i* necessary to create one cell of strain *μ*. The second assumption states that the resource uptake rates can be factored into a species-and resource-specific (but concentration independent) factor *r_μ,i_*, and a species-independent (but resource and concentration-specific) function 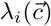. For example, 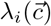 could denote the uptake rate of a pathway that imports resource *i*, while *r_μ,i_* denotes the constitutive expression of that pathway in an individual of strain *μ*. In this picture, strains can differ in their overall expression of a given pathway, but have limited ability to tune its biochemical properties.

We assume that the resource fluxes and concentrations are both large, such that the dilution and noise terms can be neglected in Eq. (A2). Following Ref. (32), we also assume a separation of timescales between the dynamics of resource concentrations, such that the resource concentrations reach a quasi-equilibrium 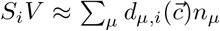 before the strain abundances start to change significantly. Under these assumptions, we can eliminate the concentration variables entirely, and obtain a set of coarse-grained dynamics for the strain abundances:

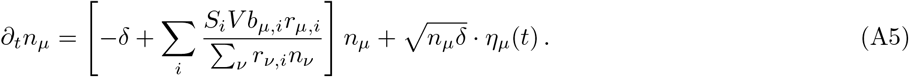

In this model, the dynamics of the total number of individuals, 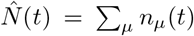, does not close, due to the *μ* dependence in the biomass conversion factor *b_μ,i_*:

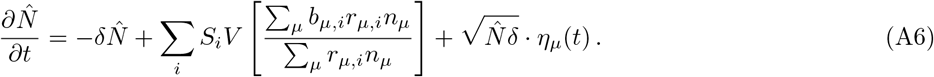

However, if we assume that *b_μ,i_* ≈ *b_i_* (similar to our previous assumption that 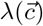 is independent of *μ*), then the equation for 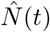 closes, and we find that 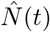 rapidly approaches a steady-state value 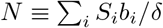 on a timescale of order 1/*δ*, with fluctuations of order 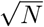. Such fluctuations become irrelevant in the large *N* limit, which suggests that we rewrite the dynamics in terms of the strain frequencies, 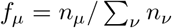. Following the derivation in Ref. (46), the dynamics of the frequencies *f_μ_* satisfy

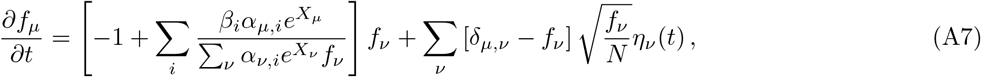

where we have defined the normalized parameters

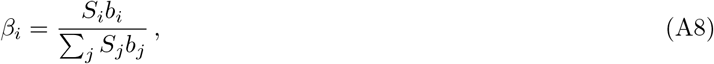

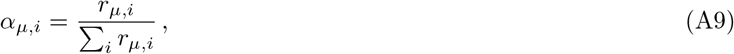

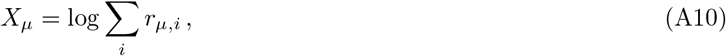

and time is measured in units of *δ*^−1^.

### 2. Subdivided environment model

The familiar form of Eq. (A7) suggests that these dynamics can also be obtained from a generalization of the standard Wright-Fisher model, in which the population is periodically subdivided into separate environments. In this model, the strains in environment *i* produce a number of gametes proportional to their Wrightian fitness, *W_μ,i_*. After a period of growth, *Nβ_i_* gametes are chosen from each environment and mixed together to obtain the next generation. The expected fraction of individuals in the next generation is

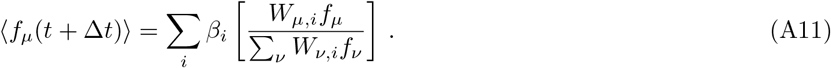

When 〈*f_μ_*(*t* + Δ*t*) – *f_μ_*(*t*)〉 is small, this update rule has the same continuum limit as Eq. (A7), with

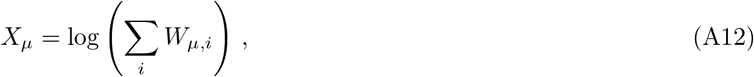

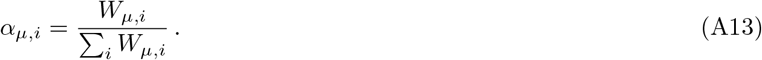

### 3. Deterministic Lyapunov function

The deterministic dynamics possess a Lyapunov function

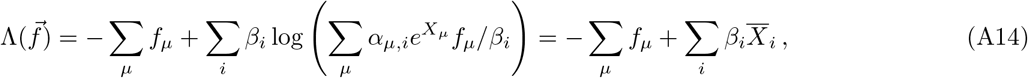

for which

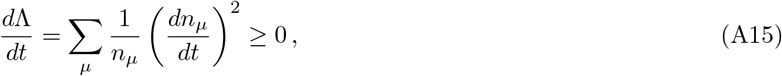

and which is convex and bounded from above. Among other things, this implies that the deterministic dynamics have a unique equilibrium that is approached at long times. We exploit this fact in the simulations in Appendix E.

## Appendix B: Competition for two resources

In this section, we derive our main results for the two resource case. The major advantage of this limit is that the multidimensional resource space reduces to the scalar interval (0,1). Without loss of generality, we will write everything in terms of the first resource component, defining *β* = *β*_1_ and *α*_*μ*,1_ = *α_μ_*, with the remaining components *β*_2_ = 1 − *β* and *α*_*μ*,2_ = 1 − *α_μ_* fixed by the normalization condition. Following the description in the main text, we will begin by analyzing the competition between two strains, and then consider the effects of adding a third strain to a pair of previously coexisting strains.

### 1. Competition between two strains

To anayze the competition between two strains, we let *α*_1_ and *α*_2_ denote the strategy vectors of the two strains, and let Δ*X* = *X*_2_ − *X*_1_ denote the fitness difference between them. We arbitrarily designate strain 1 as the “wildtype” and consider the frequency of the “mutant strain”, *f* ≡ *f*_2_. With these definitions, Eq. (1) can be rewritten in the form,

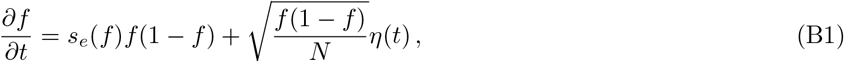

which is familiar from single-locus population genetics (47), where we have defined an effective frequency-dependent selection coefficient,

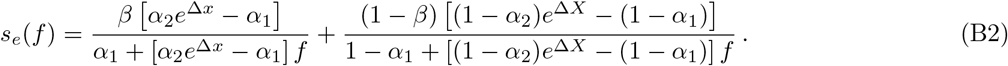

Our main results can be derived from limiting versions of this basic model.

#### Invasion of a new strain

The invasion of a new strain corresponds to the *f* → 0 limit, in which Eq. (B1) reduces to the linearized form,

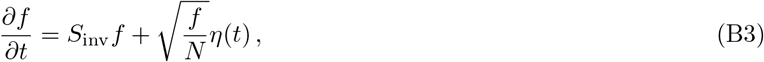

with an invasion fitness *S*_inv_ defined by

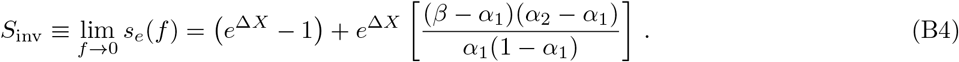

Eq. (B3) can be solved using standard methods (48), so we will simply quote the relevant results here. For initial frequencies small compared to the 1/*NS*_inv_, the genetic drift term dominates, and there is a high probability that the mutant will drift to extinction. However, with probability *p*_est_ = 2*NS*_inv_*f*(0), the mutant will drift to frequency ~1/*NS*_inv_, after which point the selection term dominates over genetic drift. This “established” lineage will then grow deterministically as 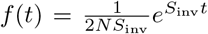, which can be matched onto the full nonlinear (but deterministic) solution as *f* increases further. The full solution is somewhat unwieldy, but the first-order nature of the ODE shows that *f*(*t*) cannot decrease as *t* → ∞. Thus, once the mutant establishes, the deterministic dynamics will never drive the mutant close enough to the drift barrier that extinction becomes likely again. This suggests that the branching process description will be valid as long as *f*(*t*) remains sufficiently small during the duration of the establishment process that *f*(*t*) ≪ 1 and *s_e_*(*f*) ≈ *S*_inv_. This will be true provided that these conditions are satisfied at the drift barrier, 1/*NS*_inv_, which leads to the conditions

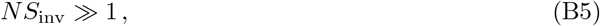

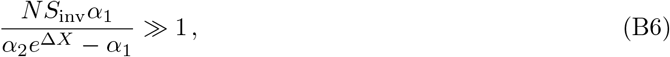

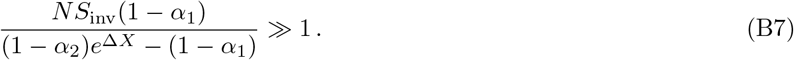

These conditions can be satisfied simultaneously for sufficiently large *N*.

#### Stable coexistence

If *S*_inv_ > 0 and the mutant is lucky enough to establish, then the frequency-dependent selection term will either drive the mutant to fixation (*f* = 1) or else stabilize at some intermediate frequency *f*^*^. As described in the main text, stable coexistence requires that the reciprocal invasion fitness,

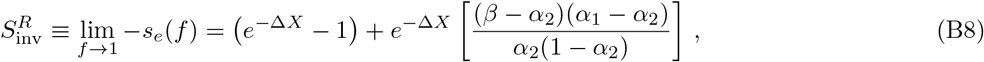

is also positive. Solving this equation when 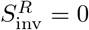 yields the critical fitness threshold

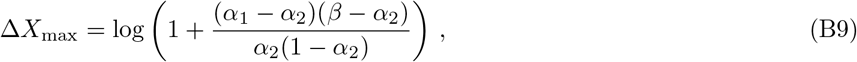

which reduces to Eq. (7) in the main text in the near-ESS limit. We might naively assume that this threshold would be equivalent to the fitness that gives strain 2 a higher uptake rate on *both* individual resources, i.e. *α*_2_*e*^Δ*X*^ ≥ *α*_1_ and (1 – *α*_2_)*e*^Δ*X*^ ≥ 1 – *α*_1_. Although this is indeed a sufficient condition for strain 2 to fix, the true thresholds in Eqs. (7) and (B9) are much weaker conditions, which depend on the environmental supply vector *β*. This means that in practice, stable coexistence will be disrupted long before one of the strains is uniformly better than the other.

When the conditions for stable coexistence are met, the equilibrium frequency *f*\* is obtained from the condition that *s_e_*(*f*\*) = 0. From Eq. (B2), we see that this can only happen if *α*_2_*e*^Δ*X*^ – *α*_1_ and (1 – *α*_2_)*e*^Δ*X*^ – (1 – *α*_1_) have different signs, i.e. neither strain is uniformly better than the other. Solving for *f*\*, we find that

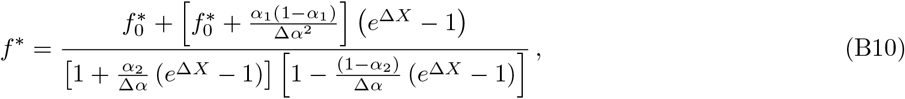

where 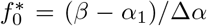 is the equilibrium frequency in the absence of any fitness differences. When *f* = *f*\*, the resource-specific mean fitnesses 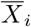 take on the values

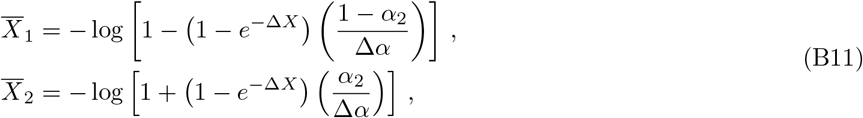

which are independent of the resource supply vector *β*. This extends the “environmental shielding” behavior derived in the neutral limit by Ref. (32): when two strains coexist on two substitutable resources, the strain frequencies evolve so that the remaining selection pressures take on values that are independent of the environment, and depend only on the identities of the coexisting strains. We will revisit this behavior again in the multi-resource case below.

In the limit that fitness differences are small [specifically, when Δ*X* is small compared to 1, *α*_2_/Δ*α*, and (1–*α*_2_)/Δ*α*], Eq. (B11) reduces to the linearized version,

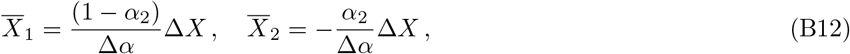

while Eq. (B10) reduces to the linear relation quoted in Eq. (8) in the main text. This defines a second fitness scale,

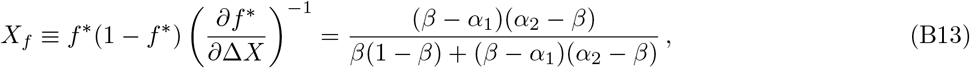

over which *f*\*(Δ*X*) changes significantly. Note that *X_f_* has approximately the same scaling behavior for small and large *β* – *α* as the critical threshold Δ*X*_max_ in Eq. (7).

For frequencies close to *f*\*, the selection term again grows small compared to the genetic drift term. Linearizing Eq. (B1) around *f* ≈ *f*\*, the fluctuations *δf* = *f* – *f*\* are described by

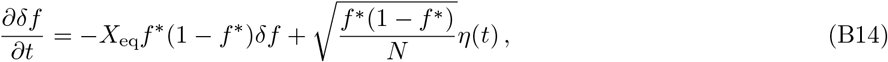

where we have defined the equilibrium restoring fitness

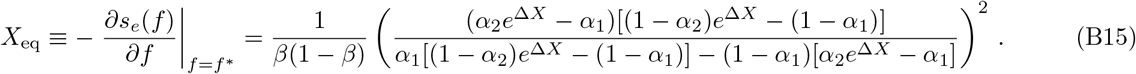

In the limit that Δ*X* ≪ 1, this becomes

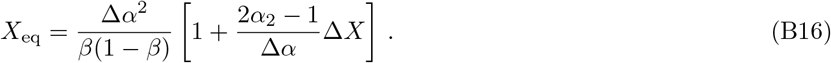

This model can again be solved using standard methods (49). The stationary distribution of *δf* tends toward a normal distribution with mean zero and standard deviation 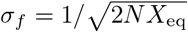, which decays on a timescale ~1/*X*_eq_*f*\*(1 – *f*\*). The quasi-deterministic model is therefore self-consistent provided that

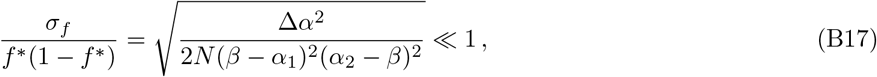

which can be satisfied for sufficiently large *N*.

The fluctuations in *f* lead to similar fluctuations in the resource-specific mean fitnesses, 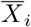, whose first order contribution is given by

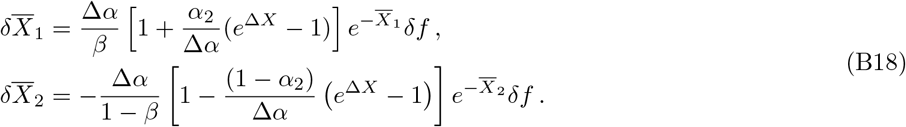

### 2. Competition between three strains

Having characterized the dynamics for a pair of strains, we next consider a scenario in which a third strain is introduced into a stable ecosystem where a pair of strains already coexist. Without loss of generality, we will assume that the third strain is a mutant version of the second strain, with fitness *X*_3_ = *X*_2_ + *s* and strategy vector *α*_3_. For small initial frequencies, the mutant strain has an invasion fitness

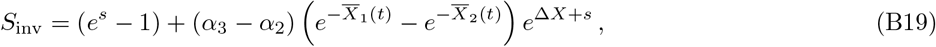

where the resource-specific mean fitnesses 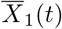 and 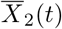 are dictated by the two strain process in Eq. (B1). We consider the implications of this expression in various special cases below.

#### No fitness differences

In a completly neutral scenario (Δ*X* = *s* = 0), the resource-specific mean fitnesses are solely determined by the fluctuations 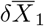 and 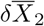 from Eq. (B18), and Eq. (B19) reduces to

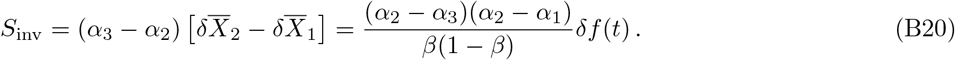

Since 〈*δf*(*t*)〉 = 0, this agrees with the deterministic results of Ref. (32), who found that all further invasion fitnesses vanish in a neutral population when the ecosystem is fully expoited. However, our stochastic analysis shows that fluctuations can induce momentary selection pressures of order

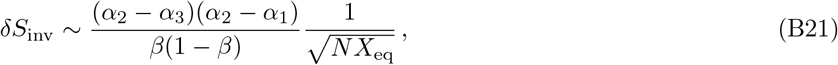

which can be large compared to 1/*N*. However, these momentary selection pressures average out to zero over a timescale 1/*X*_eq_*f*\*(1 – *f*\*). When *N* is large, this is much shorter than the timescale ~1/*δS*_inv_ required for the mutant lineage to escape the drift barrier. This shows that internal fluctuations cannot induce anomalous establishment events in our model. To leading order in *N*, ecological selection pressures vanish in a neutral population when two strains coexist on two substitutable resources.

#### Pure fitness mutations

In the case where the mutant lineage is created by a pure fitness mutation, *α*_3_ = *α*_2_, and the invasion fitness reduces to

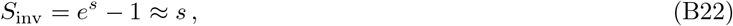

which is identical to the standard Wright-Fisher model. This justifies our interpretation of *X_μ_* as a fitness parameter. Eq. (B22) is a slightly stronger result, since it implies that general fitness mutations continue to establish at the same rate, regardless of the structure of the ecosystem. When such a mutation establishes, it is guaranteed to displace its parent strain, resulting in a two-strain competition between strain 3 and strain 1, which now differ in fitness by an amount Δ*X* + *s*. If Δ*X* + *s* ≥ Δ*X*_max_ from Eq. (7), then stable coexistence will be disrupted, and strain 3 will take over the entire population. On the other hand, if Δ*X* + *s* < Δ*X*_max_, the mutant will only displace its parent strain, and will be prevented from sweeping through the entire population. Instead, the successful mutation will shift the equilibrium frequency by an amount

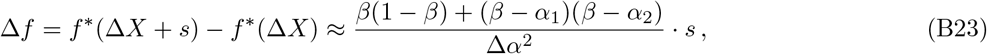

where we have employed the linearized approximation for *f*\* from Eq. (8) in the main text.

#### Pure strategy mutations

If the mutant lineage is created by a pure strategy mutation (*s* = 0), then the invasion fitness reduces to

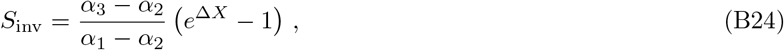

where we have retained only the leading order contribution as *N* → ∞. The Δ*X* ≪ 1 limit is listed in Eq. (13) in the main text. The interpretation of this expression, and the various scenarios that can arise after establishment, are described in the main text as well.

### 3. Evolution of a single-locus ecology

The results above allow us to analyze the effects of further evolution in our consumer resource model. As a first pass, we focus on a simplified scenario, in which strategy mutations switch between two fixed strategy vectors, *α*_1_ and *α*_2_, and occur at rate *U_α_*. We assume that *α*_1_ and *α*_2_ span *β*, so that the strains can stably coexist. We also assume that *α*_1_ and *α*_2_ are sufficiently close to *β* that we can invoke the near-ESS limits of various expressions above. We note that while this assumption is also employed in the adaptive dynamics literature (22, 42), our model also differs from these results in a key way, as it includes *α* that go beyond the infinitesimal evolution assumption in adaptive dynamics.

Our model also differs from the canonical adaptive dynamics scenario in that it includes pure fitness mutations, which occur at rate *U_XρX_*(*s*). We assume that the tails of *ρ_X_*(*s*) are sufficiently light that the distribution can be approximated by a characteristic beneficial fitness effect (50), which we will also denote by the generic variable *s* below. Our analysis here will focus on the strong-selection weak mutation (SSWM) regime that arises in the limit that *N* → ∞ and *U_α_* + *U_X_* → 0. The first assumption guarantees that genetic drift is only relevant when mutations are sufficiently rare, so that the establishment process can be modeled by the branching process techniques above. The second assumption guarantees that all mutations establish or go extinct before the next mutation occurs, so that they can be described by the two- and three-strain competition processes above. Violations of this assumption are considered in more detail in a following section.

#### No strategy mutations

We first consider the dynamics under pure fitness mutations when *U_α_*/*U_X_* = 0. We assume that the population has just diversified into a pair of coexisting strains with fitness difference Δ*X* = 0, and equilibrium frequency 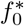. Pure fitness mutations will occur in each clade at rate *NU_X_f*\* and *NU_X_*(1 – *f*\*), respectively. According to Eq. (B22), these establish with probability *p*_est_ = 2*s*, sweep through their parent clade, and result in a new fitness differential,

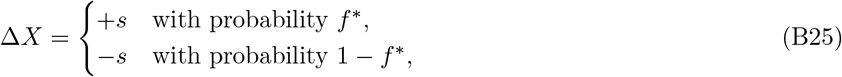

which depends on the genetic background in which the mutation arose. This fitness differential will lead to a shift in the equilibrium frequency Δ*f* = ±*s*/*s_c_* described by Eq. (9) in the main text. If Δ*f* < –*f*\* or Δ*f* > 1 – *f*\*, then stable coexistence will be disrupted, and the mutant strain will take over the entire population. This will occur whenever

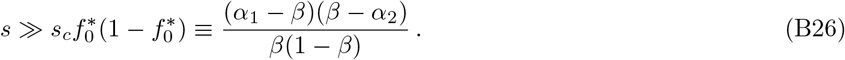

In this regime, the lifetime of coexistence is of order *τ*_collapse_ ~ 1/*NU_X_ s* (the time that it takes for one fitness mutation to occur).

In the opposite regime, when 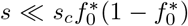, individual fitness mutations lead to small shifts in *f*\*, and many such mutations must accumulate before stable coexistence is disrupted. In this case, we can model the changing equilibrium frequency using an effective diffusion process. In an interval of time *δt*, the fitness differential changes by *δ*Δ*X* = *s*(*k*_2_ – *k*_1_), where *k*_2_ and *k*_1_ denote the number of fitness mutations that accumulate in the *f*\* and 1 – *f*\* backgrounds, respectively. In the weak mutation limit, these occur as a Poisson process with rates 2*NU_X_sf*\**δt* and 2*NU_X_s*(1 – *f*\*)*δt*, respectively, so that

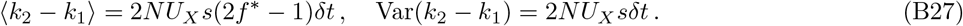

The fitness difference Δ*X* can therefore be described by an effective diffusion process,

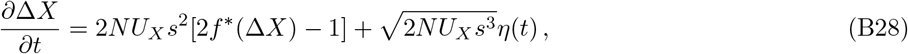

where the equilibrium frequency *f*\* itself depends on Δ*X* through Eq. (8) in the main text. Changing variables from Δ*X* to *f*\*, we obtain Eq. (10) in the main text. For our detailed calculations, it will be somewhat more convenient to work with the rescaled variables *Y* = 2*f*\* – 1 and *k* = 2*NU_X_st*, which yields a related equation

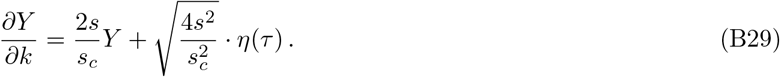

This is similar to the equation for the drift-induced fluctuations in Eq. (B14), except that the bias is now a destabilizing force, rather than a stabilizing one. This reflects the fact that larger clades are more likely to acquire beneficial mutations in the weak mutation limit, which leads to further increases in frequency. We can quantify the strength of this snowballing effect by analyzing the ultimate fixation probability of strain 1 (i.e., the probability that *Y* → 1) as a function of the current value of *Y* = 2*f*\* – 1. Eq. (B29) implies a corresponding backward equation for the fixation probability

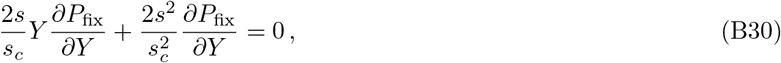

whose solution is given by

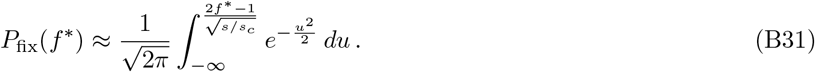

This function undergoes a sharp transition near 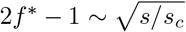. When 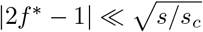, fixation and extinction of the clade are equally likely, while for 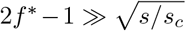, fixation is virtually guaranteed. This transition has a simple interpretation in terms of the relative strengths of the bias and noise terms in Eq. (10): 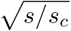 represents a critical frequency difference above which the bias dominates over the noise term. Since 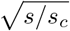 is itself a small parameter in the *s* ≪ *s_c_* regime, this implies that the random portion of the clade competition process is confined to frequencies near 50%. Reversals from frequencies near *f*\* ≈ 0 or *f*\* ≈ 1 are asymptotically unlikely.

To investigate the dynamics of this process, we analyze the mean squared frequency difference 〈*Y*^2^〉. Using Eq. (B29), we can derive a closed moment equation for 〈*Y*^2^〉

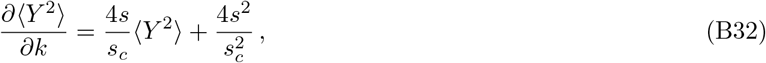

whose solution is given by

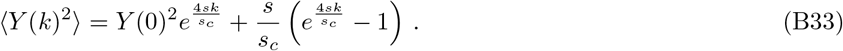

Solving for *k* and converting back to units of time, we find that

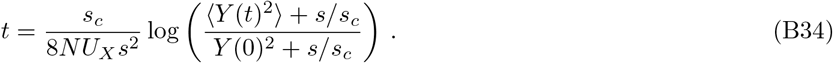

The behavior of this function has a simple heuristic interpretation based on the fundamental timescales of Eq. (10). Starting from 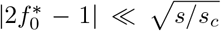, the clade frequencies will wander diffusively for a time 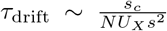 until the frequency difference reaches 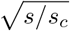, after which point the major clade deterministically acquires mutations for 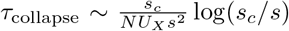 more generations until it reaches fixation. On the other hand, if the clades start with a frequency difference 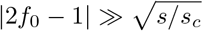, then the major clade will deterministically fix within 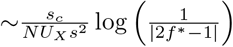 generations

##### Including strategy mutations

We can use the results above to analyze the case where *U_α_*/*U_X_* > 0. For very low values of *U_α_*/*U_X_*, strategy mutations will rarely occur before the ecosystem collapses according to the process described above. In this case, the main effect of strategy mutations is to re-diversify a population that consists of a single ecotype. The invasion fitness of such a mutation is therefore given by Eq. (3) in the main text, and will vary depending on which ecotype dominates the population.

We can therefore distinguish between two regimes. If 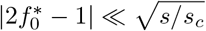, then both ecotypes are equally likely to fix, and the average invasion fitness is

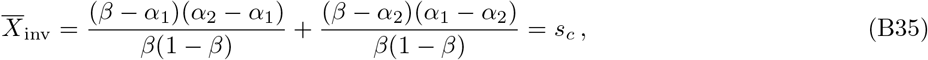

This leads to a diversification timescale *τ*_diversify_ ~ 1/*NU_α_s_c_*, and the diversification-selection balance in Eq. (11) in the main text. On the other hand, if 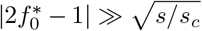 then the clade with the larger initial frequency will typically be the one that fixes. Without loss of generality, we will relabel the strains so that *f*\* always represents this clade. In this scenario, the average invasion fitness is instead given by 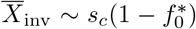, which is strictly smaller than *s_c_*. In this case, the diversification-selection balance is given by

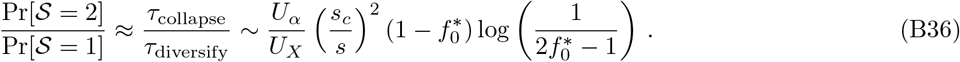

For still larger values of *U_α_*/*U_X_*, strategy mutations will start to occur before one of the clades has fixed in the population. If the mutation occurs in the fitter clade, it will have an invasion fitness *S*_inv_ = |Δ*X*|, and will reset the fitness difference to zero if it establishes. On the other hand, if the mutation occurs in the less fit clade, it will have a negative invasion fitness and will not be able to establish. Thus, the net effect of these strategy mutations is to set Δ*X* = 0 at a time-dependent rate

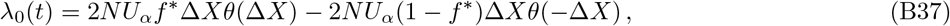

where *θ*(*z*) is the Heaviside step function, and *f*\*(*t*) and Δ*X*(*t*) are determined by the effective diffusion process in Eq. (10) in the main text. The first successful strategy mutation will occur on a characteristic invasion timescale determined by the implicit relation

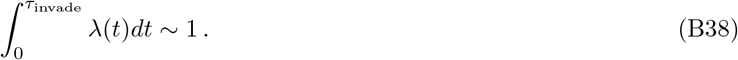

Since the fitter strain will typically be the most abundant as well, Eq. (B37) will only differ by a factor of two from the much simpler expression

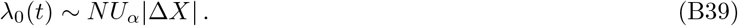

Since Eq. (B38) is only accurate up to an order one factor, we will use this simpler approximation for λ_0_(*t*) instead.

Based on these definitions, we can obtain a self-consistent solution to Eq. (B38) in various regimes. If *U_α_* ≫ *U_X_*, then strategy mutations will arise much faster than individual mutations. In this case, a lucky fitness mutation will establish in one of the clades after a time of order 1/*NU_X_s*, so that

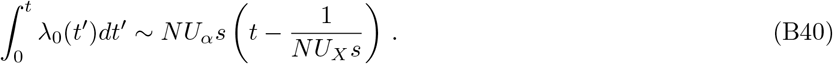

This yields an invasion timescale

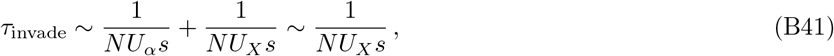

which is self-consistent provided that *U_α_* ≫ *U_X_*.

If *ρ*_invade_ ≫ 1/*NU_X_ s*, then multiple fitness mutations will accumulate before the first successful strategy mutation arises. If *τ*_invade_ ≪ *τ*_drift_ then the fitness differential Δ*X* wanders diffusively as 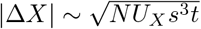, and

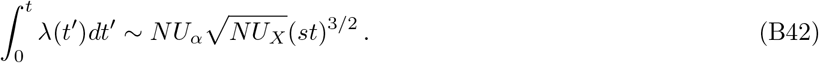

This leads to an invasion timescale of order

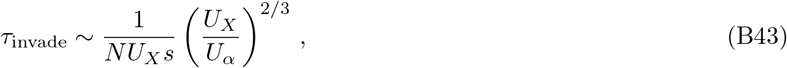

which is self consistent provided that *U_α_* ≪ *U_X_* ≪ *U_α_* (*s_c_*/*s*)^3/2^.

If *τ*_invade_ ≫ *τ*_drift_, or if the initial frequency differential already exceeds the critical value 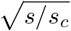, then the successful strategy mutation will occur when Δ*X* is growing deterministically as

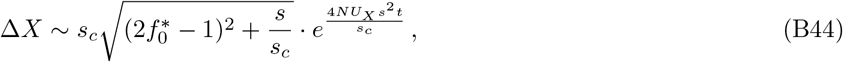

so that

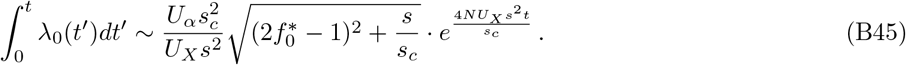

If *τ*_invade_ ≪ *τ*_collapse_, this leads to an invasion timescale,

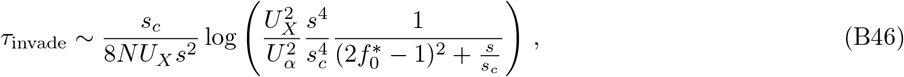

which will be self-consistent provided that 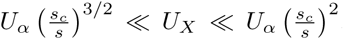. Finally, for 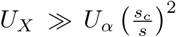, strategy mutants are sufficiently rare that the ecosystem will typically collapse and re-diversify before invasion can occur. In this case, *τ*_invade_ formally diverges. The various regimes for *τ*_invade_ are summarized in Eq. (12) in the main text.

##### The effects of clonal interference

In our analysis above, we have focused on the weak mutation limit, in which only two or three strains exist within the population at any one time. While this enabled many analytical simplifications, it is also known that many microbial populations lie outside this regime. This is particularly true for many microbial evolution experiments in which stable coexistence has been observed to evolve spontaneously. While a thorough analysis of this regime is beyond the scope of the present work, we will summarize the key differences that are likely to arise in the effective diffusion process in Eq. (10) in the main text.

Outside of the weak-mutation limit, many established beneficial mutations will be driven to extinction due to clonal interference with other beneficial mutations that happen to segregate at the same time (51). In the limit that clonal interference is strong (*NU_X_* ≫ 1), this has two main consequences. First, the rate of adaptive substitution scales much more weakly with *N* than the linear expectation *NU_X_s* from the SSWM limit. In the case of coexisting strains, this will also apply to the subpopulations *Nf*\* and *N*(1 – *f*\*) that correspond to the two clades. As a result, the bias term in Eq. (10) will be significantly reduced (and essentially vanishes in the limit of strong clonal interference). Second, clonal interference causes the rate of adaptation to become more deterministic in addition to reducing it, since it is no longer limited by the supply of beneficial mutations. The dynamics of these fluctuations are poorly understood in the general case, though Ref. (52) has shown that they lead to a long-term diffusion constant,

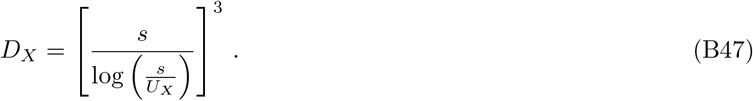

for the total fitness gain in a model similar to ours. Thus, as long as *Nf*\*(1 – *f*\*) remains sufficiently large that clonal interference within each clade remains strong, we expect the effective diffusion model in Eq. (10) to be better approximated by the limiting form

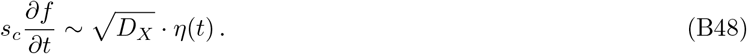

Due to the weaker bias term, we expect that the relative frequencies of the clades can undergo dramatic reversals before one or the other accumulates a fitness advantage that is large enough it to fix. Interestingly, such reversals have been observed in a long-term experiment in *E. coli* (13). However, a more thorough analysis of this clonal interference regime remains an interesting avenue for future work.

## Appendix C: Competition for many resources

In this section, we show how many of the results derived in the two-resource case can be extended to systems with larger numbers of resources. Most of these results will apply for arbitrary values of 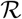, but we are particularly interested in the qualitative differences that arise in the many resource limit where 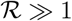.

### 1. Invasion of a mutant strain

We begin by considering a mutation that occurs in an ecosystem with an arbitrary number of coexisting strains, with equilibrium resource-specific mean fitnesses, 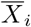. Without loss of generality, we will assume that the mutation occurs in the *μ* =1 strain, and leads to a new phenotype 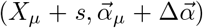, where the stragy perturbation must satisfy the normalization constraint 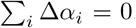. The invasion fitness for the resident strain *μ* must be zero, since it is by definition present at the ecological equilibrium. Using this fact, along with the normalization condition on 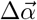, one can show that the general invasion fitness for the mutant is given by

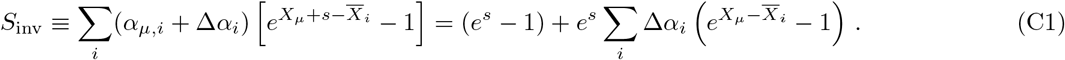

In the near-ESS limit where *s, X_μ_*, and 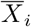 are all small compared to one, this expression reduces to Eq. (14) in the main text.

### 2. Ecological equilibria

The invasion fitness in Eq. (C1) depend on the structure of the stable ecosystem through the resource-specific mean fitnesses, 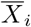, which depend on the equilibrium strain frequencies 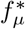 through the definition in Eq. (2). Compared to the two-resource case above, it is generally more difficult to calculate the ecological equilibrium for a set of strains when 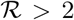. Part of this difficulty is caused by the vector nature of the resource space, which can no longer be projected down onto a single scalar dimension. However, this is more than just a book-keeping issue — there are also fundamentally new kinds of ecological equilibria that can arise when 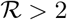. In a two-resource system, ecological equilibria are either monocultures (with 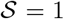 resident strains), or else contain the maximum number of coexisting strains permitted by the environment 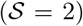. However, when 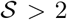, one can also have stable coexistence at any intermediate value of 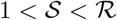, in addition to the saturated state with 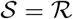. These two classes of equilibria turn out to have very different properties.

#### Saturated ecosystems

The saturated stable state 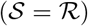 is the closest analogue of the two-resource equilibrium that we studied in Appendix B. In this case, we can obtain an explicit solution for the strain frequencies, 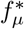, and resource-specific mean fitnesses, 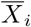, attained at equilibrium as a function of the phenotypes 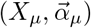 of the resident strains. By definition, the per capita growth rate (*∂_t_* log *f_μ_*) of each resident strain must vanish at equilibrium, which yields a system of 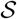 equations for the 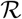 resource-specific mean fitnesses:

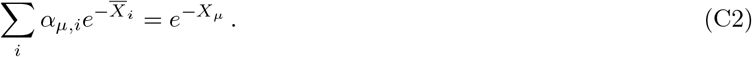

When *k* = *p*, this system can be inverted to obtain

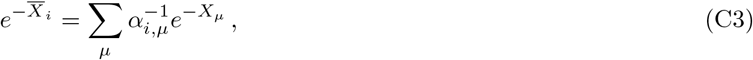

where 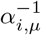 is the left inverse of *α_μ,i_*. In the limit that |*X_μ_* – *X_ν_*| ≪ 1, this reduces to Eq. (15) in the main text. Using the definition of 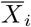 in Eq. (2) in the main text, we obtain a second system of 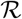 equations for the *k* equilibrium frequencies:

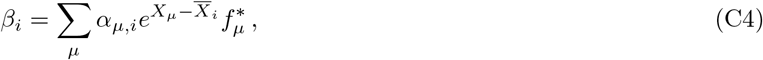

which is the non-neutral generalization of Eq. (4) in the main text. Again, when 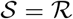, we can invert this system to obtain

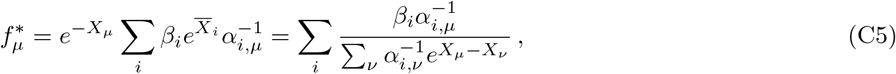

since the left and right inverses are equal in this case. In the limit that |*X_μ_* – *X_ν_*| → 0, we obtain the leading order contribution

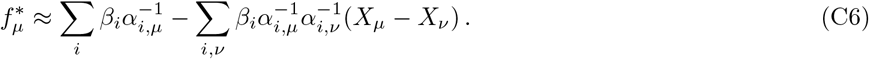

To gain intuition into these formulae, we consider a set of strains whose resource strategies are a mixture of specialist and generalist components:

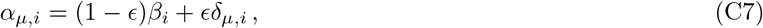

where 0 ≤ *ϵ* ≤ 1 provides a measure of the “distance” between the resource strategies. In this case, the inverse matrix has the asymptotic limits

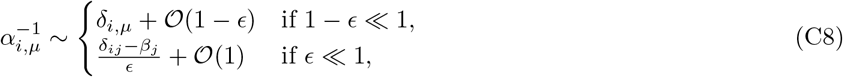

so that

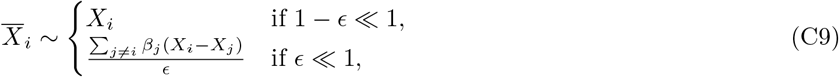

and

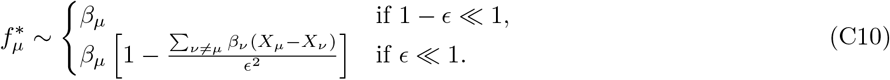

#### Unsaturated ecosystems

In contrast to the saturated case, when the number of surviving species is less than the number of resources 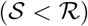 the equations in Eq. (C2) underdetermine the resource-specific mean fitnesses, 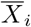, so we must invoke the non-linear constraints in Eq. (C4) to jointly solve for 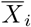 and 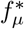. Alternatively Ref. (33) has shown that the equilibrium values of 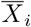 can be obtained from the solution of a convex optimization problem, subject to the constraints in Eq. (C2). In particular, if we define the transformed variable 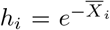, then the equilibrium value of *h_i_* is the solution to the convex optimization problem

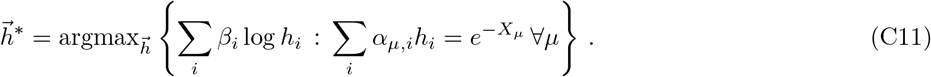

In fact, this method yields a general solution for the equilibrium value of 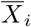 for *any* initial collection of strains, provided that the equality constraints in Eq. (C11) are replaced by inequalities (≤). Given the equilibrium values of 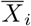, the surviving species correspond to the indices *μ* where the equality condition is satisfied. The corresponding values of 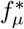 satisfy the (generally overdetermined) set of equations in Eq. (C4), which can be inverted using constrained linear regression. We employ this technique to implement the SSWM simulations in Appendix E below.

We note that since the objective function in Eq. (C11) depends on *β_i_*, the equilibrium values of 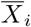 will also generally depend on the environmental supply vector in an unsaturated ecosystem, in contrast to the *β*-independent values obtained in the saturated case. Thus, the ecosystem is no longer able to dynamically adjust to “shield” the internal selection pressures from the current state of the environment (32, 33, 34). Shifts in *β_i_* can therefore lead to new opportunities for evolutionary adaptation.

### 3. Evolution in a binary resource usage model

Since there are few empirical constraints on the genetic architecture of resource strategies in the limit of many resources 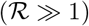, we focused on a toy “binary usage” model similar to the one considered by Ref. (33). In this model, genomes can either encode the ability to utilize a given resource or not (e.g. through the presence or absence of a particular pathway), so that the resource strategy is of the form

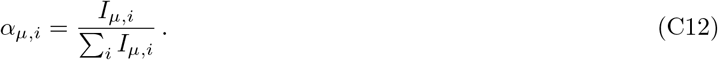

where *I_μ,i_* ∈ 0, 1 is a binary indicator variables. Individuals can acquire loss-of-function mutations at rate 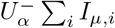, which cause one of the values of *I_μ,i_* = 1 to switch to *I_μ,i_* = 0. We also assume that they can acquire gain of function “mutations” (e.g. horizontal acquisition of a gene from the environment) at rate 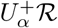, which force a randomly chosen uptake rate to the *I_μ,i_* = 1 state. Under these assumptions, the pure mutation dynamics will lead to an binomial ensemble of resource strategies, analogous to the one considered by Ref. (33), with a “success probability” of

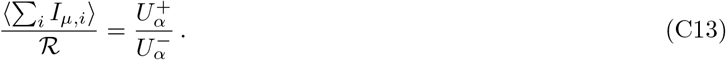

For simplicity, we will assume that the 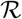 resources are all supplied at nearly identical rates. Note that in the completely symmetric case 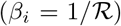, a “generalist” strain with *I_μ,i_* = 1 will constitute a marginal evolutionary stable state. To avoid this pathological behavior, we will consider small perturbations around the completely symmetric state:

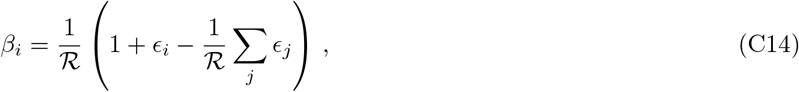

where the *ϵ_i_* are small random perturbations drawn from some distribution, and sorted in descending order 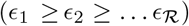. For simplicity, we will assume that the *ϵ_i_* are i.i.d. Gaussian variables with scale *ϵ* ≪ 1. The steep tail ensures that the maximum perturbation scales as

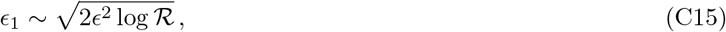

and can be bounded to be sufficiently small for a suitable choice of *ϵ*. Under these assumptions, an ecosystem comprised of a single “generalist” strain will still have nonzero ecological selection pressures encoded by the resource-specific mean fitnesses,

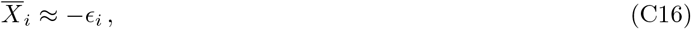

so that some alternate resource strategies will be favored to invade.

By simulating evolutionary dynamics in this model in the weak mutation limit for various values of *U_X_* and *s* (Appendix E.2), we find that the long-term structure of the ecosystem tends toward a state in which there is a single generalist strain and 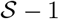 single loss-of-function variants that have recently descended from this strain (Figs. 5 and S2-S4). In the limit that *U_X_*/*U_α_* → 0, this state must also coincide with a saturated state 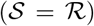. If we let *f*_1_ denote the frequency of the generalist strain, then the equilibrium frequencies are given by

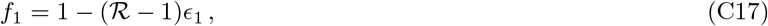

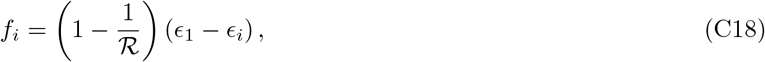

where we have assumed that 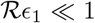. In other words, all resources except the one with the largest value of *ϵ_i_* will have a loss-of-function strain. In the limit that 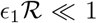, the loss-of-function strains will constitute a tiny fraction of the population, and most mutations will arise in the generalist strain. In particular, the accumulation of fitness mutations will cause the fitness of the generalist strain to grow as *X*_1_ ~ *NU_X_s*^2^*t*. We therefore wish to understand when and how this fitness differential drives some of the loss-of-function variants to extinction.

Due to the symmetry of the system, if *j* strains are driven to extinction, these must be strains with loss-of-function mutations in genes with the next *j* largest values of *ϵ_i_*, i.e. *i* = 2,…, *j* + 1. Let *X_c_*(*j*) denote the critical value of *X*_1_ required for these *j* strains to go extinct. Expanding Eq. (1) in the main text to lowest order in *X*_1_, (1 – *f*_1_), 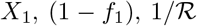, and *ϵ*, the equilibrium frequencies satisfy

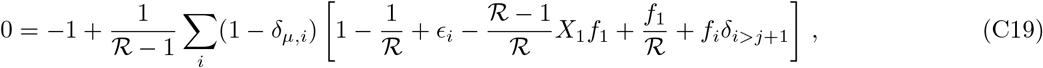

or

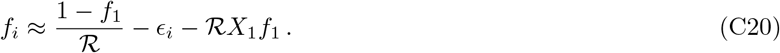

To self consistently solve for 1 – *f*_1_, we sum over 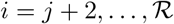 to obtain:

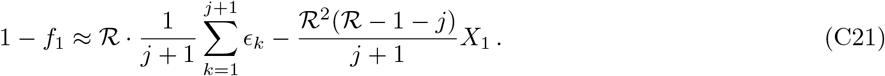

Substituting this back into our expression for *f_i_*, we obtain:

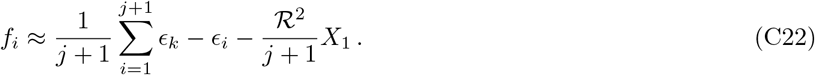

We can then self-consistently solve for *j* by setting *f*_*j*+1_ = 0. For example, if *j* = 1, then we have

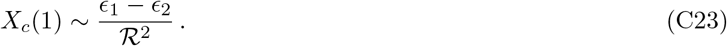

This will be a quenched random variable, since we have assumed that the *ϵ_i_* are randomly distributed. Given our Gaussian assumption, the typical value of *ϵ*_1_ – *ϵ*_2_ will occur for

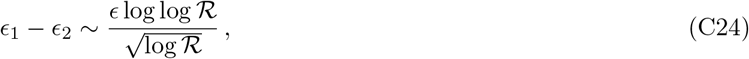

which yields

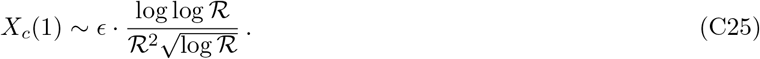

On the other hand, if 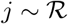, we have

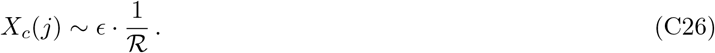

These two fitness scales are separated by a gap of order

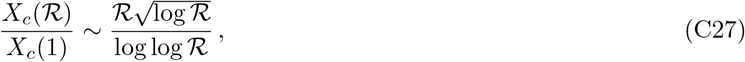

which grows increasingly large as 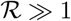.

We can use these results to derive heuristic expressions for the number of species 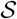 at steady state as a function of *U_α_*/*U_X_*. We first consider the limit where 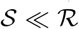 As mentioned above, the generalist strain comprises the vast majority of the population, so that to a first approximation, we can assume that all fitness and strategy mutations occur on this genetic background. Furthermore, since 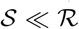, most loss of function mutations will target a resource *i* that does not already have a loss-of-function variant, where the resource-specific mean fitness is given by

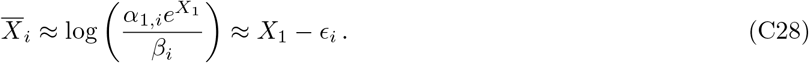

According to Eq. (14), the invasion fitness for a loss-of-function variant that targets resource *i* is given by

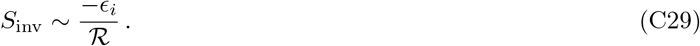

Since these loss of function mutations are produced from the generalist background at rate *NU_α_* per resource, the number of coexisting strains increases at rate

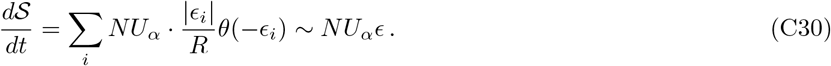

in the absence of fitness mutations.

However, as we mentioned above, the accumulation of fitness mutations will cause the fitness of the generalist strain to grow as *X*_1_(*t*) ~ *NU_X_s*^2^*t*. Since loss-of-function variants do not acquire further fitness mutations of their own, their fitness is frozen at whatever fitness the generalist strain had at the time that the mutation arose. The fitness difference between the mutant and the generalist therefore grows with time until it reaches a critical value 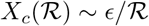, at which point the loss-of-function variant is driven to extinction. This gives rise to two characteristic dynamical regimes depending on whether 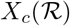 is larger or smaller than the effect *s* of a typical fitness mutation.

If *s* 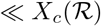, then the generalist lineage must acquire multiple fitness mutations to drive one of the loss-of-function variants to extinction. To a first approximation, the fitness difference between the generalist and the *j*th most-recently created loss-of-function variant in this regime is given by

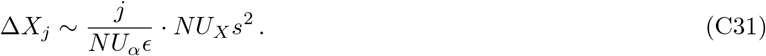

The number of coexisting ecotypes 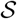 at steady-state is therefore determined by the relation 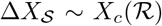, which reflects a balance between the elimination of the oldest loss-of-function variant due to the accumulation of fitness mutations and the production of new loss-of-function variants through strategy mutations. Solving for 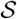, we obtain the scaling relation,

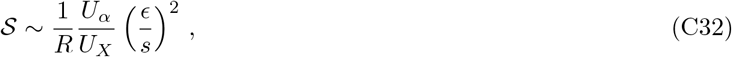

listed in Eq. (17) in the main text.

On the other hand, if 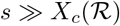, then a single fitness mutation in the generalist strain is sufficient to drive loss-of-function variants to extinction. Before this mutation arises, all the loss-of-function variants will share the same fitness difference (Δ*X_j_* = 0), and this value suddenly shifts to 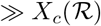 once the successful fitness mutation occurs, driving all of the existing loss-of-function variants to extinction. This leads to oscillations in 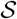 in the time between successive fitness mutations, which range from 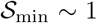 immediately after the fitness mutation arises, to a maximum value of 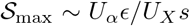 right before the next mutation arises. Since the loss-of-function variants accumulate linearly with time, this leads to a time-averaged value

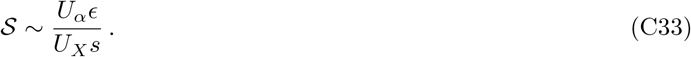

Both expressions should remain valid up to the point where there is an appreciable probability that new loss-of-function mutations target a resource that already has pre-existing variant 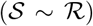. However, there can still be a broad intermediate regime between this point and the point where the ecosystem is completely saturated 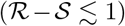. The saturated state will coincide with the evolutionary steady-state if the generalist strain is able to seed fitter loss-of-function variants into all the relevant resource dimensions before *X*_1_(*t*) increases to the point *X_c_*(1), where the first strains start to go extinct. Once again, there are two characteristic timescales depending on whether *X_c_*(1) is large or small compared to *s*.

If *s* ≪ *X_c_*(1), then the generalist lineage must acquire multiple fitness mutations to before the first loss-of-function variants are driven to extinction. This will happen over a timescale,

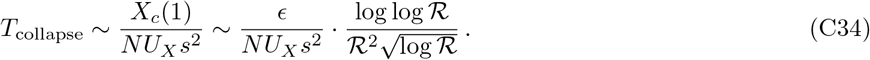

During this time, loss-of-function mutations will occur in the generalist background at rate 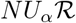 and will establish with probability ~*X*_1_(*t*). Since the loss-of-function mutations are chosen randomly, 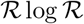 such establishments are required to cover the total number of resource dimensions with high probability (53). This requires a timescale *T*_div_ that satisfies

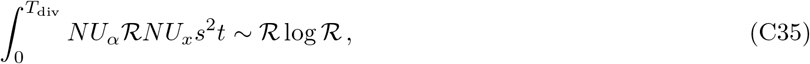

or

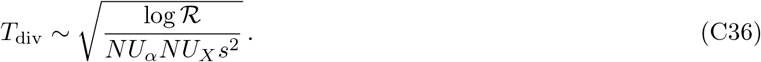

The ecosystem will remain saturated if *T*_collapse_ ≫ *T*_div_, which leads to the condition

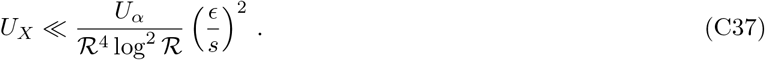

We can compare this point to the transition to 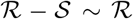 from Eq. (17), which shows that these regimes are separated by a gap of order

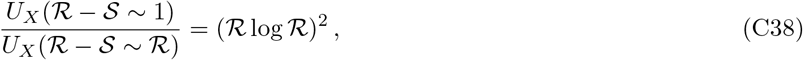

while

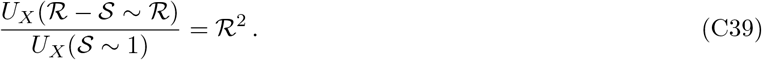

### 4. Limits on the number of utilized resources

The fragility of the diversification-selection balance in Appendix C.3 can be attributed in large part to the emergence of a fit generalist strain that is able to utilize all of the available resources. In practice, however, there might be biological constraints or other costs that limit the number of resources that a given strain can metabolize. This leads us to consider an extension of our binary resource usage model, in which the maximum number of utilized resources is capped at some value 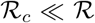. In this way, we can consider complex ecosystems 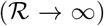 while restricting the metabolic repertoire of any given strain. A full analysis of this model is beyond the scope of the present work. Instead, we will outline a heuristic calculation that suggests that diversification-selection balance at large 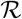 is achieved for substantially higher values of 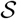 than in Appendix C.3 above.

We first note that when 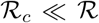, multiple strains are required to cover the available resources. The minimum possible number of strains is 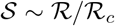, which is achieved when each of the strains specializes on a disjoint subset of 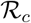 resources. To lowest order in *ϵ*, the frequencies of these strains are given by

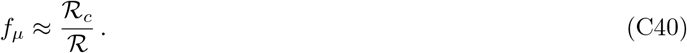

With the same genetic architecture of strategy mutations that we assumed above, this state will form the basis of the new diversification-selection balance. Generalizing our analysis above, this state will consist of 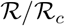 independent copies of the diversification-selection balance in Appendix C.3, except with 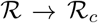. The strains that utilize 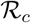 resources will be prevented from branching into new resources because of the maximum resource capacity. Meanwhile, single loss-of-function mutants on these backgrounds will be too small to acquire a gain-of-function mutation before their parent acquires enough fitness differences to drive them to extinction.

However, this behavior is strongly dependent on the specific genetic architecture that we assumed, as well as our focus on the SSWM limit. In larger populations, there may be a substantial probability for strains to acquire multiple strategy mutations in a short period of time, which would allow them to break out of their resource neighborhood. To mimic this effect in the SSWM limit, we can introduce a new rate 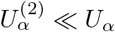 to represent the probability that two strategy mutations arise in the same lineage in a single generation. In particular, we will use this new rate to model *resource swap* events, in which one of the currently utilized resources is deleted and replaced with a randomly drawn resource. As above, we will assume that this mutation rate scales with the number of utilized resources, so that the net rate is given by 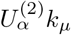, where 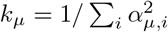.

In this augmented model, if we start from the set of 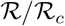 disjoint strains, then the fitnesses of these strains will wander diffusively as 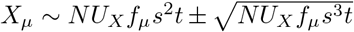, so that the typical fitness differences between a pair of strains is of order 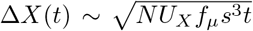. These fitness differences will create a selection pressure for strategy mutations that swap a resource from one of the fitter strains with a resource from one of the less fit strains. Such a mutation will have an invasion fitness

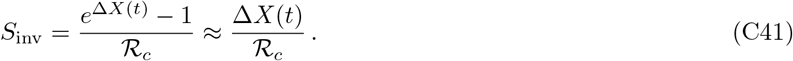

Successful swap mutations will be produced on a timescale *τ*_diversify_ that satisfies

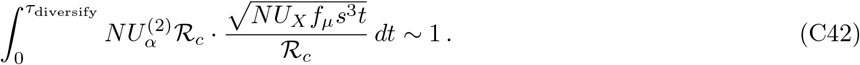

Solving for *τ*_diversify_, we obtain

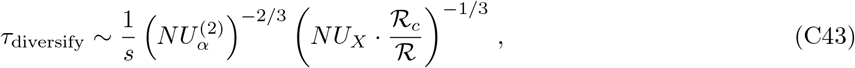

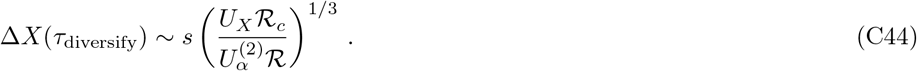

Once the successful swap mutation invades, it will create a new ecotype that coexists with the parent clade, as well as the ecotype that currently utilizes the new resource. To lowest order in 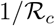, the equilibrium frequency is given by

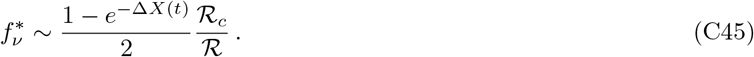

When Δ*X* ≪ 1, this frequency will be small compared to the other dominant ecotypes. As above, fitness mutations will therefore preferentially accumulate in the dominant ecotypes, causing the fitness advantage, Δ*X*(*t*), of the swap mutant to decrease over time at rate 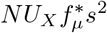. After a time of order

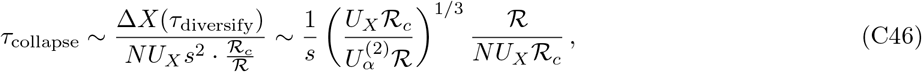

the fitness of the less fit ecotype will have caught up to the swap mutant, and the latter will be driven to extinction. The ratio between *τ*_collapse_ and *τ*_diversify_ is therefore given by

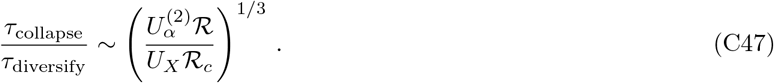

If this ratio is sufficiently large, then new swap mutants will typically establish before the fitness differences drive any of the existing swap mutants to extinction. For fixed 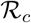, this will become increasingly true 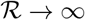.

On the other hand, if *τ*_collapse_ ≪ *τ*_diversify_, then a typical resource swap mutation will be driven to extinction before the next arises. However, because the timing of the swap mutations is a random process, anomalously late mutations may occur for which 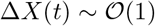. In this case, the swap mutant is no longer rare compared to its parent, and there is strong selection pressure for loss- (and later gain-) of-function mutations to arise in this mutant background.

Together, these arguments suggest that the simplest generalization of the steady-state in Appendix C.3 will generally be unstable whenever we impose a cap on the number of utilized resources, and that the corresponding diversification-balance will be attained for much higher values of 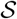 than we would expect based on our previous analysis. Further analysis of these dynamics are left for future work.

## Appendix D: Connections to adaptive dynamics

Our model shares certain features associated with the traditional models studied in adaptive dynamics (22, 42), though it also differs from these models in several key ways. In this section, we attempt to make this connection more explicit, using the notation and terminology employed in the adaptive dynamics literature. As adaptive dynamics relies on the weak mutation limit, we will confine our discussion to this regime as well.

### Two resources, no fitness differences

For simplicity, we will start by considering the two-resource case in the absence of fitness differences, where individuals are described by a scalar resource phenotype *α*. To make the connection with adaptive dynamics explicit, we will define a rescaled trait,

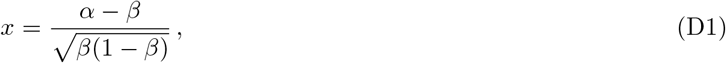

such that *x* → 0 as *α* → *β*. Following Ref. (22), we then let *s_x_*(*y*) denote the invasion fitness of a mutant of phenotype *y* in a monomorphic population of phenotype *x*. In the neighborhood of *x* → 0, Eq. (3) shows that *s_x_*(*y*) takes on a simple quadratic form

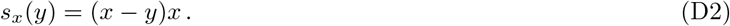

Under the standard adaptive dynamics assumption that *y* is infinitesimally close to *x*, the phenotypes will evolve in the direction of the fitness gradient,

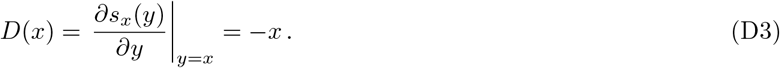

This gradient vanishes for *x* = 0 which shows that *x*\* =0 (or *α*=*β*) is an evolutionarily singular strategy. This strategy is *convergence stable*, in that infinitesimal mutations drive the population toward *x* = 0 when |*x*| > 0. However, it is only marginally ESS-stable, since *∂*^2^*s_x_*(*y*)/*∂y*^2^ = 0 at *x* = 0. The second derivative classification in Ref. (22) also shows that stable dimorphisms can coexist in the neighborhood of *x*\*.

These two features combine to make our evolutionarily singular strategy behave as *both* an evolutionarily stable strategy (ESS) and an evolutionary branching point. On the one hand, *x*\* =0 resembles an ESS because no mutant strains are favored to invade once the population reaches *x*\*. On the other hand, *x*\* =0 resembles an evolutionary branching point because the population will typically branch into a stable dimorhpism once *x* – *x*\* approaches the typical spacing between mutants. Thus, in practice, the population will always branch before it reaches the ESS, even if this is excluded under truly infinitesimal evolution. However, unlike a traditional branching point where 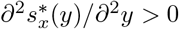, there is no further selection to drive the branched phenotypes *x*_1_ and *x*_2_ away from each other once branching has occurred. We showed in the text that this can be viewed as a generic feature of a saturated ecosystem (where 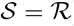) when there are no general fitness differences between strains.

#### Resource continuum, no fitness differences

One might ask why the evolutionarily singular strategy is so peculiar in our model, given that consumer-resource theory is often touted as an example of evolutionary branching points in the adaptive dynamics literature (26). The key difference is that in this existing literature, the trait *x* does not usually refer to the uptake rate of a single resource, but instead is used to parameterize an entire curve of resource uptake rates for a continuum of different resources. To choose a simple example, one might imagine that the resources denote seeds of different sizes, which are indexed by a continuous parameter *z*. The function *β*(*z*) then represents the distribution of seed sizes supplied by the environment, which is often assumed to have a Gaussian form

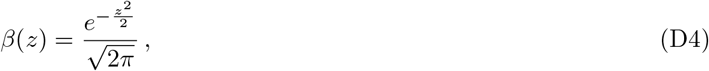

centered at some special value *z* = 0. Individual uptake rates are often assumed to have a similar Gaussian shape

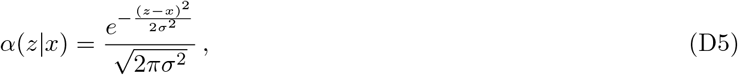

with a preferred value of *z* = *x* and a characteristic width *σ*. The trait *x* is then subject to further evolution, rather than the individual uptake rates *α*(*z*). Substituting these functions into Eq. (1) (with *X_μ_* = 0), the invasion fitness for phenotype *y* in a monomorphic population of phenotype *x* is given by

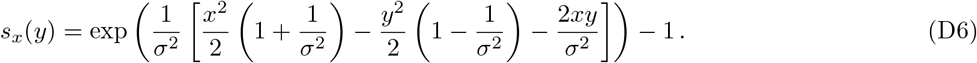

The fitness gradient *∂s_x_*(*y*)/*∂y* vanishes when *x* = 0, showing that *x*\* = 0 is an evolutionarily singular strategy, as anticipated. The second derivative is given by

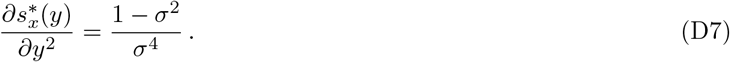

For *σ* > 1, *x*\* is a true ESS, while for *σ* < 1, *x*\* is a true evolutionary branching point. In the neighborhood of *x*\*, selection will act to drive the phenotypes further apart from each other after branching has occurred.

We can understand this behavior using the intuition developed in the main text. Since there are an infinite number of resources in this model, the ecosystem is certainly not saturated when 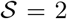. Thus, we can expect much of the selection pressure to focus on bringing the population averaged uptake rate 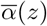 closer to the environmental supply rate *β*(*z*). When the niche width *σ* is larger than the range of resources supplied by the environment, the best way to do this is with a single strain centered at *x* = 0. Branching is therefore not favored. On the other hand, if *σ* is smaller than the range of supplied resources, then the ecosystem as a whole can match the environmental supply rate better if there are two strains centered at intermediate locations on the real axis (|*x* – *y*| > 0).

In this way, we see that the 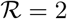 resource case, far from being pathological, serves as a basic building block that allows us to understand more complex scenarios that are often considered in the literature. It also illustrates how the genetic architecture of the uptake rates (in this case, whether the *α*(*z*) can evolve independently or are restricted to the Gaussian family) can play a key role in determining the emergent dynamics of the model.

#### Directional selection as an intermediate asymptotic

We now return to the two-resource case above and examine how changes in the general fitness (*X*) alter the adaptive dynamics analogy. Individuals are now described by a two-dimensional phenotype, (*X*, *α*). Generalizing our analysis above, we will now define a two-dimensional trait space:

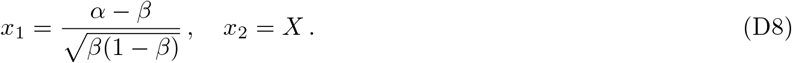

In this notation, the invasion fitness in Eq. (5) becomes

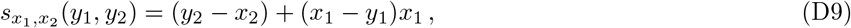

whose fitness gradient is given by

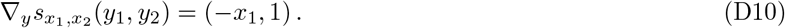

As expected, the general fitness dimension always selects for phenotypes that increase *X*, regardless of the value of *α*. As a consequence, there are no longer any evolutionarily singular strategies in this model, so the formal classification such points in the adaptive dynamics framework does not apply any more. Nevertheless, we have seen that behaviors very similar to evolutionary branching still occur in our model if we project down onto the *x*_1_ coordinate. Furthermore, the old evolutionarily singular strategy at 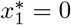 continues to play a key role in these dynamics. The major difference is that these ecologically stable polymorphisms are now only quasi-stable under evolutionary perturbations, as our analysis in the main text shows that further fitness evolution can drive one of the ecotypes to extinction (Fig. 3B). This behavior is consistent with observations from laboratory evolution experiments (13).

**FIG. S1.**
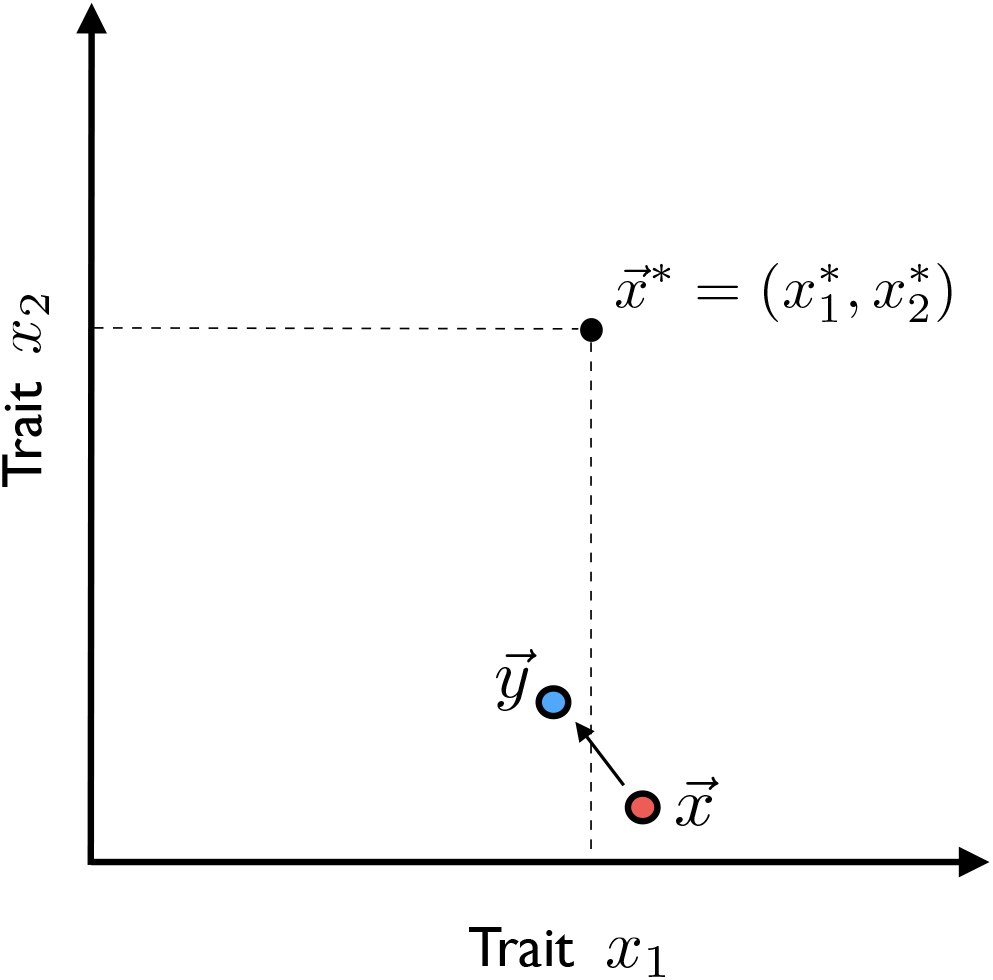
An intermediate asymptotic of adaptive dynamics. In a multidimensional phenotype space, a population that is far from the evolutionarily singular strategy can display the quasi-stable branching behavior analyzed in the main text if one of the trait dimensions (*x*_1_) is close to the singular coordinate 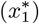. In the specific context of our consumer resource model, *x*_1_ corresponds to the resource uptake strategy (*α*), while *x*_2_ corresponds to the general fitness (*X*).

Although we have motivated this behavior with the abstract notion of “general fitness,” our analysis suggests that similar behavior will generically arise in multi-dimensional phenotype spaces whenever one of the traits (*i*) approaches its marginal branching point 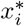, while at least one of the other traits (*j*) remains far from 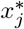 (Fig. S1). Previous work has shown that such highly assymetric approaches to a stationary point are a common feature of gradient descent dynamics in high dimensional spaces (54). In this way, our results can be viewed as an intermediate asymptotic that describes the process of ecological diversification during the asymptotically long times required for a high-dimensional trait space to approach a true evolutionarily singular strategy. A more detailed description of this regime is left for future work.

## Appendix E: Simulations

### 1. Individual-based simulations

The simulations in Fig. 1 were carried out using an individual-based, discrete generation algorithm similar to the one employed in Refs. (37, 50) for a single resource. Each simulation starts with a clonal population of *N* individuals, and in each subsequent generation, the population undergoes a selection step followed by a mutation step. At each step, we keep track of the number of individuals *n_μ_* with a given strategy vector *α_μ,i_* and general fitness *X_μ_*.

In the selection step, each lineage *n_μ_* is assigned a new size from a Poisson distribution with mean

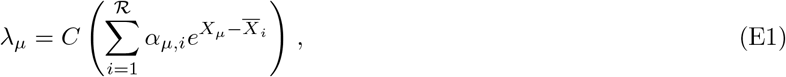

where

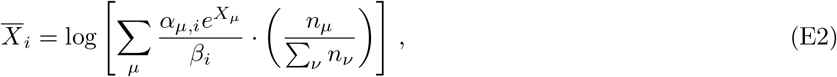

and 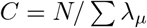 is a normalization constant chosen to ensure that the total population size remains near 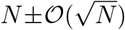.

In the mutation step, the new lineage size is pruned into multiple sublineages representing different mutations that occur on the original lineage background. With probability *U_X_*, an individual founds a new sublineage *ν* is founded with fitness *X_ν_* = *X_μ_* + *s*, where *s* is drawn from the distribution of fitness effects *ρ_X_*(*s*). With probability *U_α_*, an individual founds a new sublineage with a strategy vector 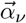 drawn from the distribution 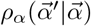.

The simulations in Fig. 1 were carried out for 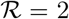 with *β* = 0.5. We utilized a Gaussian distribution of fitness effects, 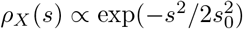, for the pure fitness mutations. The distribution of strategy mutations, *ρ_α_*(*α*’|*α*), was taken to be a beta distribution with mean *α* and coefficient of variation Var(*α*’)/*E*(*α*’)^2^ = 0.05, but with *α*’ rounded to the nearest value of 1/5, …, 4/5. The initial resource strategy for the ancestral population was chosen uniformly at random from these discrete values.

Each simulation was performed for a total of 60, 000 generations. Every 500 generations, we simulated a round of “metagenomic sequencing”. We calculated the population frequencies of all mutations present in the population, and reported these values after binomial resampling at a depth of *D* = 1000.

A copy of our implementation in `C++` is available on Github (https://github.com/benjaminhgood/consumer_resource_simulations).

### 2. SSWM simulations

To simulate the long-term dynamics of the binary usage model in Fig. 4 (Appendix C.3), we use an optimized simulation algorithm that is specifically designed for the strong-selection, weak mutation (SSWM) regime. Similar to traditional SSWM algorithms in population genetics (55), this algorithm gains an efficiency advantage by simulating only successful invasion events. In our case, however, the successful invasion events can now lead to non-trivial ecological equilibria, in addition to simple fixation.

Our simulations start with a collection of strains 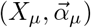 at time *t* = 0. To assess convergence to diversification-selection balance, we performed simulations for two initial conditions: (i) a single generalist strain with 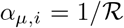 (ii) a collection of 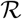 specialist strains with *α_μ,i_* = *δ_μ,i_* and *X_μ_* drawn from a Gaussian distribution with variance *σ* = 10^-7^. Figs S3 and S4 show a comparison of these two initial conditions for 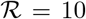. Since the agreement is generally good, we utilized the more rapidly converging generalist initial conditions for the main simulations in Figs 4 and S2.

After drawing the initial condition, we first calculate the ecological equilibrium, 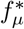, for this collection of strains using the convex optimization procedure in Appendix C.2, using the MOSEK software package (56). This algorithm yields the equilibrium values of 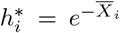 and the set of ecotypes 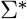 that survive at equilibrium. Within this subset, the equilibrium frequencies are obtained from the solution of the linear system in Eq. (C4), which will be overdetermined when 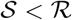. We obtain a solution to this system by solving the constrained least squares problem,

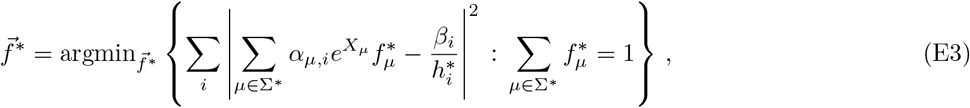

using the SciPy library (57).

Once the initial ecological equilibrium is obtained, the simulation proceeds via a series of virtual timesteps, each of which represents the successful invasion of a single mutation. In each step, we first enumerate the set of fitness and strategy mutants that are generated from mutations on each of the current strains *μ*, and calculate their corresponding invasion fitness from Eq. (C1). We use these values to calculate the net rate of successful invasions from each mutation type. We assume that fitness mutations confer a characteristic fitness benefit *s*, so that the rate of successful fitness mutations in strain *μ* is given by

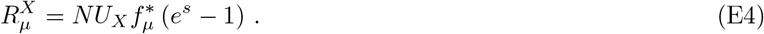

The rate of successful loss-of-function mutations for resource *i* is given by

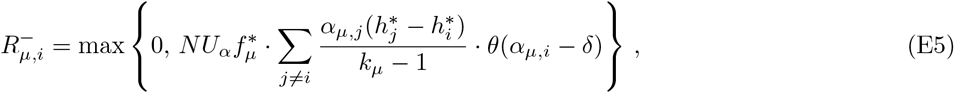

where 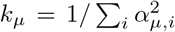 is the current number of resources utilized by strain *μ*, *θ*(*z*) is the Heavisde step function, and *δ* is an infinitesimally small positive number so that the step function is well-defined. The rate of successful gain-of-function mutations is given by an analogous expression,

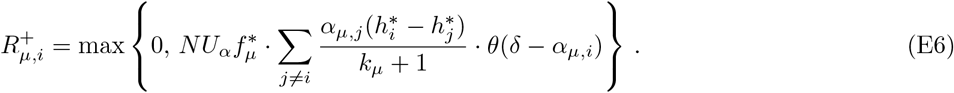

Since these successful invasion events arise as a compound Poisson process, the time *T*_est_ to the next successful invasion event is exponentially distributed with rate

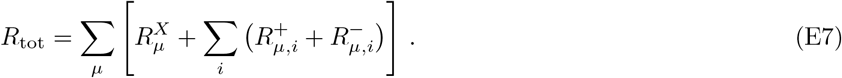

Using the Poisson thinning property, the identity of the invading mutation is chosen at random from the enumerated list with probability proportional to its corresponding *R*-value. Once the identity of the invading strain is determined, we find the new ecological equilibrium 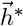 and 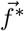 using the constrained procedure above. By assumption, the time to reach this new equilibrium is negligible compared to *T*_est_ in the SSWM limit. The current time *t* is then incremented by *T*_est_, and the process repeats itself.

We repeated this process for a total of *M* successful invasion steps until the ecosystem converged to diversification-selection balance (*M* ~ 100, 000). The simulations in Figs. 5 and S2-S4 were carried out for *ϵ* = 10^-3^ and *s* = 10^-7^, scanning through different values of *U_X_*/*U_α_*.

A copy of our implementation in Python is available on Github (http://github.com/StephenMartis/consumer-resource-many-resources).

**FIG. S2.**
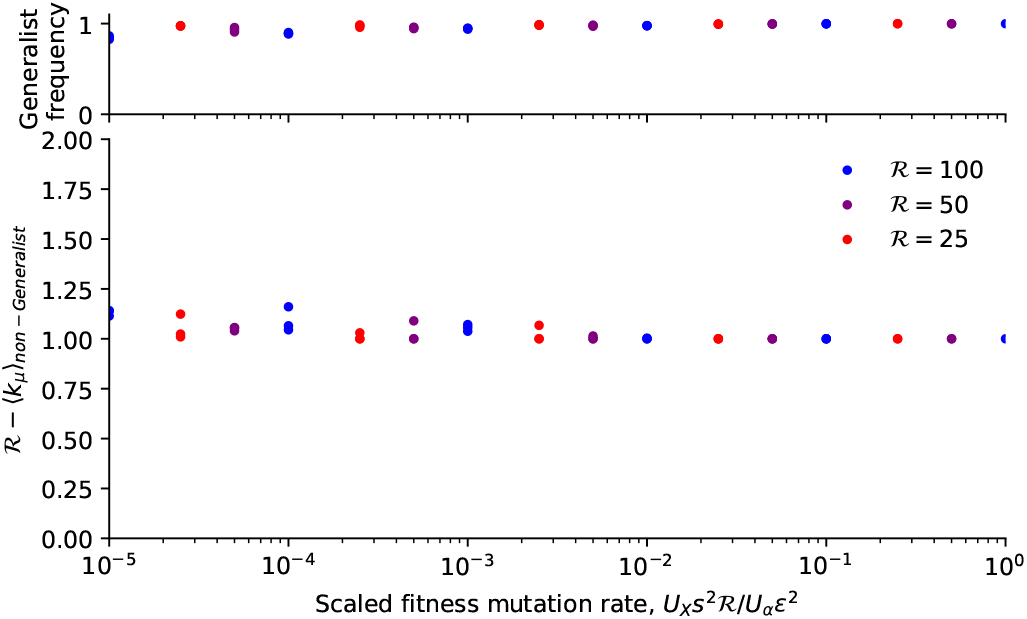
Ecological structure at diversification-selection balance in a binary usage model. (Top) For each of the simulated populations in Fig. 4, the fraction of the population occupied by the generalist ecotype, 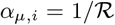. (Bottom) For the same populations, the frequency-weighted average of 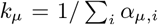 (a measure of the number of utilized resources) for the remaining non-generalist ecotypes. A value of 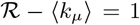 indicates that the rest of the population consists of single loss-of-function mutants that descend from the generalist background.

**FIG. S3.**
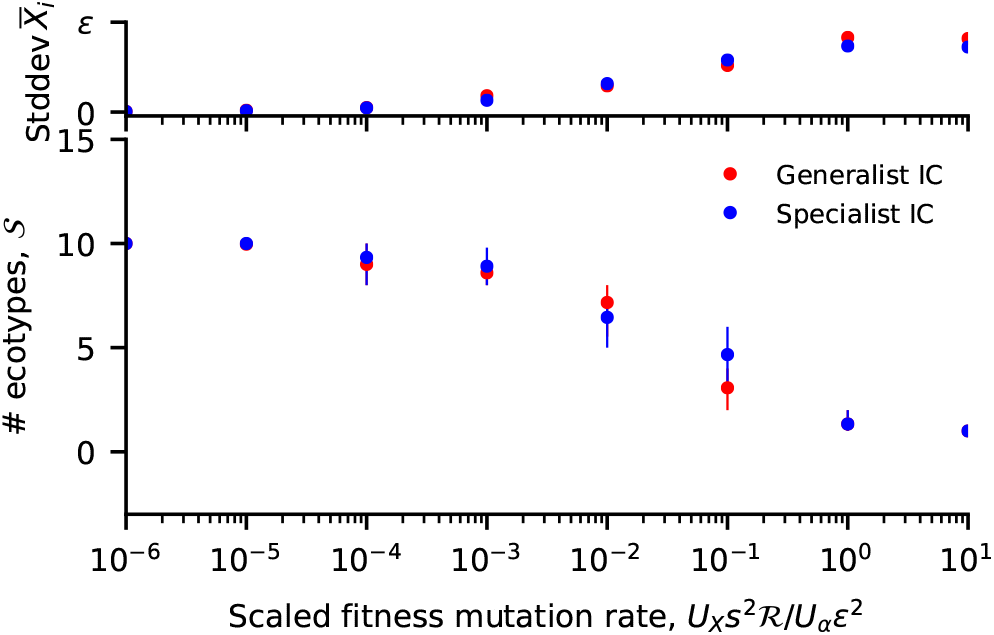
Approach to diversification-selection balance from different initial conditions. An analogous version of Fig. 4 comparing specialist and generalist initial conditions for 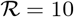.

**FIG. S4.**
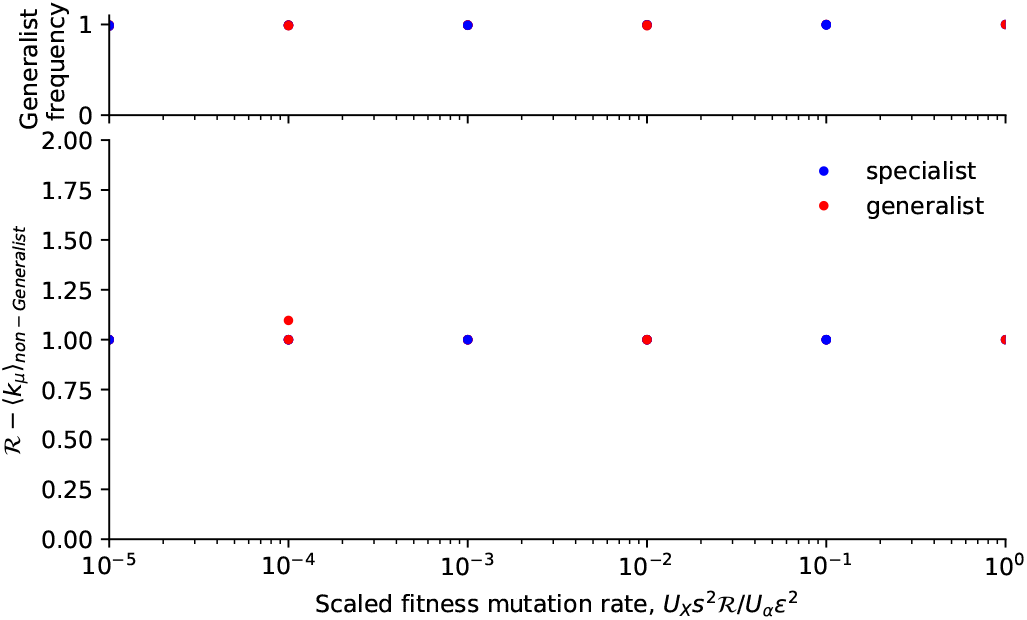
Long-term ecological structure from different initial conditions. An analogous version of Fig. S2 comparing specialist and generalist initial conditions for 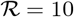.

## REFERENCES

[1] Smith JM (1983) The genetics of stasis and punctuation. Annual review of genetics 17(1):11–25.

[2] Mallet J (2008) Hybridization, ecological races and the nature of species: empirical evidence for the ease of speciation. Philosophical Transactions of the Royal Society B: Biological Sciences 363(1506):2971–2986.

[3] Nosil P, Harmon LJ, Seehausen O (2009) Ecological explanations for (incomplete) speciation. Trends in ecology & evolution 24(3):145–156.

[4] Shapiro BJ, Leducq JB, Mallet J (2016) What is speciation? PLoS genetics 12(3):e1005860.

[5] Rozen DE, Lenski RE (2000) Long-term experimental evolution in *Escherichia coli*. viii. dynamics of a balanced polymorphism. American Naturalist 155(1):24–35.

[6] Helling RB, Vargas CN, Adams J (1987) Evolution of *Escherichia coli* during growth in a constant environment. Genetics 116(3):349–358.

[7] Rainey PB, Travisano M (1998) Adaptive radiation in a heterogeneous environment. Nature 394(6688):69–72.

[8] Friesen ML, Saxer G, Travisano M, Doebeli M (2004) Experimental evidence for sympatric ecological diversification due to frequency-dependent competition in *Escherichia coli*. Evolution 58(2):245–260.

[9] Frenkel EM, et al. (2015) Crowded growth leads to the spontaneous evolution of semistable coexistence in laboratory yeast populations. Proc Natl Acad Sci USA 112(36):11306–11311.

[10] Sousa A, et al. (2017) Recurrent reverse evolution maintains polymorphism after strong bottlenecks in commensal gut bacteria. Molecular biology and evolution 34(11):2879–2892.

[11] Herron MD, Doebeli M (2013) Parallel evolutionary dynamics of adaptive diversification in *Escherichia coli*. PLoS Biology 11(2):e1001490.

[12] Traverse CC, Mayo-Smith LM, Poltak SR, Cooper VS (2013) Tangled bank of experimentally evolved burkholderia biofilms reflects selection during chronic infections. Proc Natl Acad Sci USA 110(3):E250–E259.

[13] Good BH, McDonald MJ, Barrick JE, Lenski RE, Desai MM (2017) The dynamics of molecular evolution over 60,000 generations. Nature 551(7678):45.

[14] Finkel SE, Kolter R (1999) Evolution of microbial diversity during prolonged starvation. Proceedings of the National Academy of Sciences 96(7):4023–4027.

[15] Poltak SR, Cooper VS (2011) Ecological succession in long-term experimentally evolved biofilms produces synergistic communities. ISME J 5(3):369–378.

[16] Torsvik V, Øvreås L (2002) Microbial diversity and function in soil: from genes to ecosystems. Current opinion in microbiology 5(3):240–245.

[17] Kashtan N, et al. (2014) Single-cell genomics reveals hundreds of coexisting subpopulations in wild prochlorococcus. Science 344(6182):416–420.

[18] Foster KR, Schluter J, Coyte KZ, Rakoff-Nahoum S (2017) The evolution of the host microbiome as an ecosystem on a leash. Nature 548(7665):43.

[19] Bendall ML, et al. (2016) Genome-wide selective sweeps and gene-specific sweeps in natural bacterial populations. ISME J 10(7):1589–1601.

[20] Zhao S, et al. (2017) Adaptive evolution within the gut microbiome of individual people. bioRxiv p. 208009.

[21] Garud NR, Good BH, Hallatschek O, Pollard KS (2017) Evolutionary dynamics of bacteria in the gut microbiome within and across hosts. bioRxiv. doi: 10.1101/210955.

[22] Geritz SA, Mesze G, Metz JA, et al. (1998) Evolutionarily singular strategies and the adaptive growth and branching of the evolutionary tree. Evolutionary ecology 12(1):35–57.

[23] Dieckmann U, Marrow P, Law R (1995) Evolutionary cycling in predator-prey interactions: population dynamics and the red queen. Journal of theoretical biology 176(1):91–102.

[24] Doebeli M (2002) A model for the evolutionary dynamics of cross-feeding polymorphisms in microorganisms. Population Ecology 44(2):59–70.

[25] Kondoh M (2003) Foraging adaptation and the relationship between food-web complexity and stability. Science 299(5611):1388–1391.

[26] Ackermann M, Doebeli M (2004) Evolution of niche width and adaptive diversification. Evolution 58(12):2599–2612.

[27] Ackland G, Gallagher I (2004) Stabilization of large generalized Lotka-Volterra foodwebs by evolutionary feedback. Phys Rev Lett 93(15):158701.

[28] Shoresh N, Hegreness M, Kishony R (2008) Evolution exacerbates the paradox of the plankton. Proc Natl Acad Sci USA 105(34):12365–12369.

[29] Vetsigian K (2017) Diverse modes of eco-evolutionary dynamics in communities of antibiotic-producing microorganisms. Nature Ecology & Evolution 1:0189.

[30] Neher RA (2013) Genetic draft, selective interference, and population genetics of rapid adaptation. Annual Review of Ecology, Evolution, and Systematics 44:195–215.

[31] Tilman D (1982) Resource competition and community structure. (Princeton University Press).

[32] Posfai A, Taillefumier T, Wingreen NS (2017) Metabolic trade-offs promote diversity in a model ecosystem. Phys Rev Lett 118(2):028103.

[33] Tikhonov M, Monasson R (2017) Collective phase in resource competition in a highly diverse ecosystem. Phys Rev Lett 118(4):048103.

[34] Advani M, Bunin G, Mehta P (2018) Statistical physics of community ecology: a cavity solution to macarthurs consumer resource model. Journal of Statistical Mechanics: Theory and Experiment 2018(3):033406.

[35] Mac Arthur R (1969) Species packing, and what competition minimizes. Proceedings of the National Academy of Sciences 64(4):1369–1371.

[36] Gillespie JH (1998) Population Genetics: A concise guide. (Johns Hopkins University Press, Baltimore, MD), second edition.

[37] Good BH, Desai MM (2015) The impact of macroscopic epistasis on long-term evolutionary dynamics. Genetics 199(1):177–190.

[38] Gillespie JH (2000) Genetic drift in an infinite population: The pseudohitchhiking model. Genetics 155:909–919.

[39] McInerney JO, McNally A, O’Connell MJ (2017) Why prokaryotes have pangenomes. Nature microbiology 2(4):17040.

[40] Tikhonov M, Monasson R (2017) Innovation rather than improvement: a solvable high-dimensional model highlights the limitations of scalar fitness. Journal of Statistical Physics pp. 1–31.

[41] Van Valen L (1973) A new evolutionary law. Evol Theory 1:1–30.

[42] Doebeli M (2011) Adaptive Diversification. (Princeton University Press).

[43] McDonald MJ, Gehrig SM, Meintjes PL, Zhang XX, Rainey PB (2009) Adaptive divergence in experimental populations of *Pseudomonas fluorescens*. iv. genetic constraints guide evolutionary trajectories in a parallel adaptive radiation. Genetics 183(3):1041–1053.

[44] Plucain J, et al. (2014) Epistasis and allele specificity in the emergence of a stable polymorphism in *Escherichia coli*. Science 343(6177):1366–1369.

[45] Gardiner C (1985) Handbook of Stochastic Methods. (Springer, New York).

[46] Good BH, Desai MM (2013) Fluctuations in fitness distributions and the effects of weak linked selection on sequence evolution. Theor Pop Biol 85:86–102.

[47] Ewens WJ (2004) Mathematical Population Genetics. (Springer-Verlag, New York), second edition.

[48] Kendall DG (1948) On the generalized “birth-and-death” process. The annals of mathematical statistics 19(1):1–15.

[49] Kardar M (2007) Statistical physics of fields. (Cambridge University Press).

[50] Good BH, Rouzine IM, Balick DJ, Hallatschek O, Desai MM (2012) Distribution of fixed beneficial mutations and the rate of adaptation in asexual populations. Proc Natl Acad Sci USA 109:4950–4955.

[51] Gerrish P, Lenski R (1998) The fate of competing beneficial mutations in an asexual population. Genetica 127:127–144.

[52] Fisher DS (2013) Asexual evolution waves: fluctuations and universality. J Stat Mech 2013:P01011.

[53] Feller W (2008) An introduction to probability theory and its applications. (John Wiley & Sons) Vol. 2.

[54] Kurchan J, Laloux L (1996) Phase space geometry and slow dynamics. Journal of Physics A: Mathematical and General 29(9):1929.

[55] McCandlish DM, Stolzfus A (2014) Modeling evolution using the probability of fixation: history and implications. Quarterly Review of Biology 89:225–252.

[56] Andersen ED, Andersen KD (2000) The mosek interior point optimizer for linear programming: an implementation of the homogeneous algorithm in High performance optimization. (Springer), pp. 197–232.

[57] Jones E, Oliphant T, Peterson P, et al. (2001–) SciPy: Open source scientific tools for Python.

